# Integrative multi-omics framework identifies phenotypically impactful driver pathways in glioblastoma multiforme

**DOI:** 10.1101/2025.02.28.640792

**Authors:** Junhua Zhang

## Abstract

Glioblastoma multiforme (GBM), a highly aggressive brain tumor characterized by molecular heterogeneity, demands integrative approaches to uncover driver pathways and therapeutic targets. Here, we present **gtePIDP**, a computational framework that systematically integrates multi-omics data to identify Phenotypically Impactful Driver Pathways. By evaluating mutation patterns (coverage/exclusivity) and downstream regulatory cascades, this model maps somatic alterations to phenotypic outcomes using TCGA mutation profiles, expression data, and regulatory networks (transcription factors, microRNAs). Through iterative pruning of candidate driver genes and quantifying resultant perturbations in gene expression and regulatory activities, gtePIDP prioritizes gene sets maximizing mutational significance and phenotypic impact. Our analysis uncovered pivotal GBM driver pathways involving *TP53, CDKN2A, MDM2*, and *RB1*, which disrupt key cancer-associated regulators (e.g., oncogenic: *NFATC2, MIR-370*; tumor-suppressive: *MIR-506, MIR-9, FOXJ2*), driving dysregulation of angiogenesis, apoptosis, and proliferation. Notably, we identified therapeutic axes such as *TP53/MIR185/VEGFA* and *MDM2/SMAD4/VEGFA*, suggesting anti-angiogenic targeting strategies. Furthermore, a PI3K-Akt signaling regulatory module revealed actionable targets (*CDKN2A/CREBBP/OSMR* and *TP53/MIR-9/FGF12*) for intervention. By bridging mutational landscapes with transcriptional/epigenetic dysregulation, gtePIDP advances mechanistic driver pathway discovery and prioritizes targets with clinical relevance. This study highlights the power of multi-omics integration to unravel GBM pathogenesis and accelerate precision oncology strategies. Additionally, the gtePIDP framework is readily extendable to other malignancies (such as breast carcinoma (BRCA), ovarian cancer (OV), lung adenocarcinoma (LUAD), etc.), providing a universal computational strategy for pan-cancer biomarker discovery.

## 1 Introduction

Glioblastoma multiforme (GBM) is the most aggressive and prevalent form of primary brain tumor, characterized by its high heterogeneity and poor prognosis [1]. The complexity of GBM pathogenesis underscores the need for a comprehensive understanding of the molecular mechanisms driving tumorigenesis and progression.

Cancer initiation and progression are driven by somatic alterations that disrupt key cellular pathways, conferring selective growth advantages to tumor cells. Identifying these driver pathways (or driver gene sets) and their functional consequences remains a central challenge in cancer genomics.

Traditional approaches, such as MutSig and Dendrix, focused on detecting genes with high mutation rates or mutually exclusive patterns by solving maximum weight submatrix problems (MWSP) [2–5]. Various other methods integrated two or more kinds of data and intended to reduce the search space of feasible solutions for MWSP or to increase the ability for predicting driver genes, such as the network-based approaches [6], multi-omics integration methods [7], and machine/deep learning models [8–10]. However, existing approaches mainly overlooked the downstream phenotypic impacts of mutations, such as dysregulated gene expression and altered activities of transcriptional regulators – critical factors in tumorigenesis.

Recent AI-driven advancements in cancer research demonstrate potential for biomarker discovery and personalized therapy [11, 12]. However, limitations persist, including data quality dependency, model interpretability (“black box” issues), and challenges in standardizing multi-modal inputs. In GBM, key pathways like *PI3K/AKT/mTOR* and *p53* signalings are known drivers [13, 14], but the interplay between genetic alterations, transcriptional rewiring, and transcriptional/post-transcriptional regulation by transcription factors (TFs) or microRNAs (miRNAs) remains poorly resolved.

To address these challenges, we present **gtePIDP** (genomic-transcriptomic-epigenetic Phenotypically Impactful Driver Pathways), a computational framework that integrates somatic mutations, gene expressions, and regulatory networks to identify driver pathways or gene sets with maximal phenotypic impact. Here the regulatory networks include not only those for transcriptional regulation (TFs to target genes), but also for epigenetic miRNA regulation (miRNAs to target genes, post-transcriptional regulation). A key innovation of gtePIDP lies in its dual evaluation of mutation patterns and downstream regulatory cascades. In gtePIDP, we extend the maximum weight submatrix framework [3] by incorporating gene expression data and regulatory network analysis. Specifically, we introduce an extended affinity regression (EAR) model to quantify how somatic mutations influence TF and miRNA activities, which subsequently propagate to dysregulate downstream genes. By iteratively removing candidate driver genes and evaluating resultant perturbations in expression and regulator activities – quantified via Stouffer’s Z-scores – our model identifies gene sets that maximize both mutational weight (*W(S)*) and phenotypic impact (*Z(S)*).

Applied to GBM, gtePIDP uncovered critical driver pathways involving *TP53, CDKN2A, MDM2*, and *RB1*, which perturb key cancer-associated regulators (e.g., oncogenic: *NFATC2, MIR370*; tumor-suppressive: *MIR-506, MIR-9, FOXJ2*), driving dysregulation of angiogenesis, apoptosis, and proliferation. Notably, we identified novel regulatory axes, including *TP53/MIR-185/VEGFA* and *MDM2/SMAD4/VEGFA*, suggesting targets for anti-angiogenic therapy. A PI3K-Akt signaling module revealed therapeutically actionable axes (*CDKN2A/CREBBP/OSMR(/EGFR)* and *TP53(/CDKN2A/RB1)/MIR-9/FGF12*), while *CDKN2A/CREBBP/OSMR(/EGFR)* emerged as a mediator of mesenchymal transition and therapy resistance [15, 16]. Further, *MIR-9* downregulation – driven by *TP53, CDKN2A*, and *RB1* alterations – silenced *FGF12*, a potential PI3K-Akt inhibitor, highlighting miRNA dysregulation as a hallmark of GBM pathogenesis.

In summary, this work advances cancer genomics by bridging mutational patterns with transcriptional and post-transcriptional regulatory consequences. By prioritizing driver pathways that exert maximal phenotypic impact, our framework offers a robust tool for identifying therapeutic targets in complex malignancies like GBM. Furthermore, the gtePIDP framework is readily extendable to other malignancies such as breast carcinoma (BRCA), ovarian cancer (OV), lung adenocarcinoma (LUAD), etc. The integration of multi-omics data and regulatory networks not only refines driver gene prediction but also unveils mechanistic insights into tumorigenesis, paving the way for precision oncology strategies.

## 2 Methods and materials

For a driver pathway or gene set, on one hand, it must have large coverage and exclusivity; on the other hand, it must have heavy impact on the phenotype of samples. Synthesizing these factors, we propose a novel approach for driver pathway identification.

### 2.1 Large coverage and exclusivity

We adopt the maximum weight submatrix problem to describe large coverage and exclusivity [3]. Given a binary mutation matrix *M* with *m* rows (samples) and *n* columns (genes), Vandin *et al.* introduced a weight function *W*, which is designed to find a submatrix *S* of size *m × k* in matrix *M* by maximizing this function [3]:

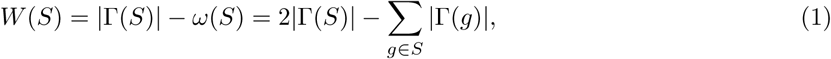

where Γ(*g*) = {*i* : *A_ig_* = 1} denotes the set of samples in which the gene *g* is mutated, Γ(*S*) = *∪_g∈S_*Γ(*g*) measures the coverage of *S*, and *ω*(*S*) = Σ*_g∈S_ |*Γ(*g*)*| − |*Γ(*S*)*|* measures the coverage overlap of *S*. Eq. 1 means to reward coverage while penalize overlap to get high exclusivity. In this way we can get a gene set with *k* genes having large coverage and exclusivity based on cancer somatic mutation data.

### 2.2 Linking somatic alterations with gene expression and activities of TFs and miRNAs

Here we employ affinity regression (AR) model [17] to investigate the impact of somatic alterations on gene expression and activities of TFs and miRNAs. Suppose we have *r* TFs, *s* miRNAs and *t* expressed genes. Given a mutation matrix *M* (*m × n*), a gene expression matrix *E*(*t × m*), and targeting relation matrices of TFs on genes *D*_1_(*t × r*) and miRNAs on genes *D*_2_(*t × s*), we can establish the following relationship:

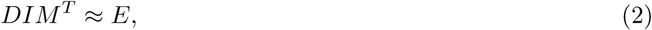

where *D* = [*D*_1_*, D*_2_], *M ^T^* is the transposition of *M*, and *I* represents the impact matrix ((*r* + *s*) *× n*) of somatic alterations on TFs and miRNAs. It is precisely *I* that we are going to learn through Eq. 2.

We introduce Eq. 2 based on a primary idea, i.e., driver pathways should heavily impact phenotypes of samples, such as gene expressions, while these expressions are directly regulated by TFs and miRNAs. In fact, Eq. 2 can also be deduced by the AR model, where the authors designed the model to learn an interaction matrix between upstream signal-transduction proteins and downstream TFs.

Similar to the AR model, Eq. 2 can be used to learn the impact matrix *I* by the mutation matrix *M*, the TF and miRNA targeting matrix *D*, and the gene expression matrix *E*. Here we call Eq. 2 as extended affinity regression (EAR) model, and we solve it by the package AR run from [17]. Once we have learned *I*, we can calculate the TFs’ and miRNAs’ activities *A*(*m ×* (*r* + *s*)):

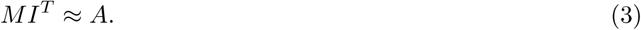

### 2.3 An integrated method for driver pathway identification

We develop a computational framework to identify driver pathways in the intent that the detected gene set not only has large coverage and exclusivity in mutations, but also it disturbs genes’ expressions and TFs’ and miRNAs’ activities as heavily as possible (Fig. 1).

**Fig. 1.**
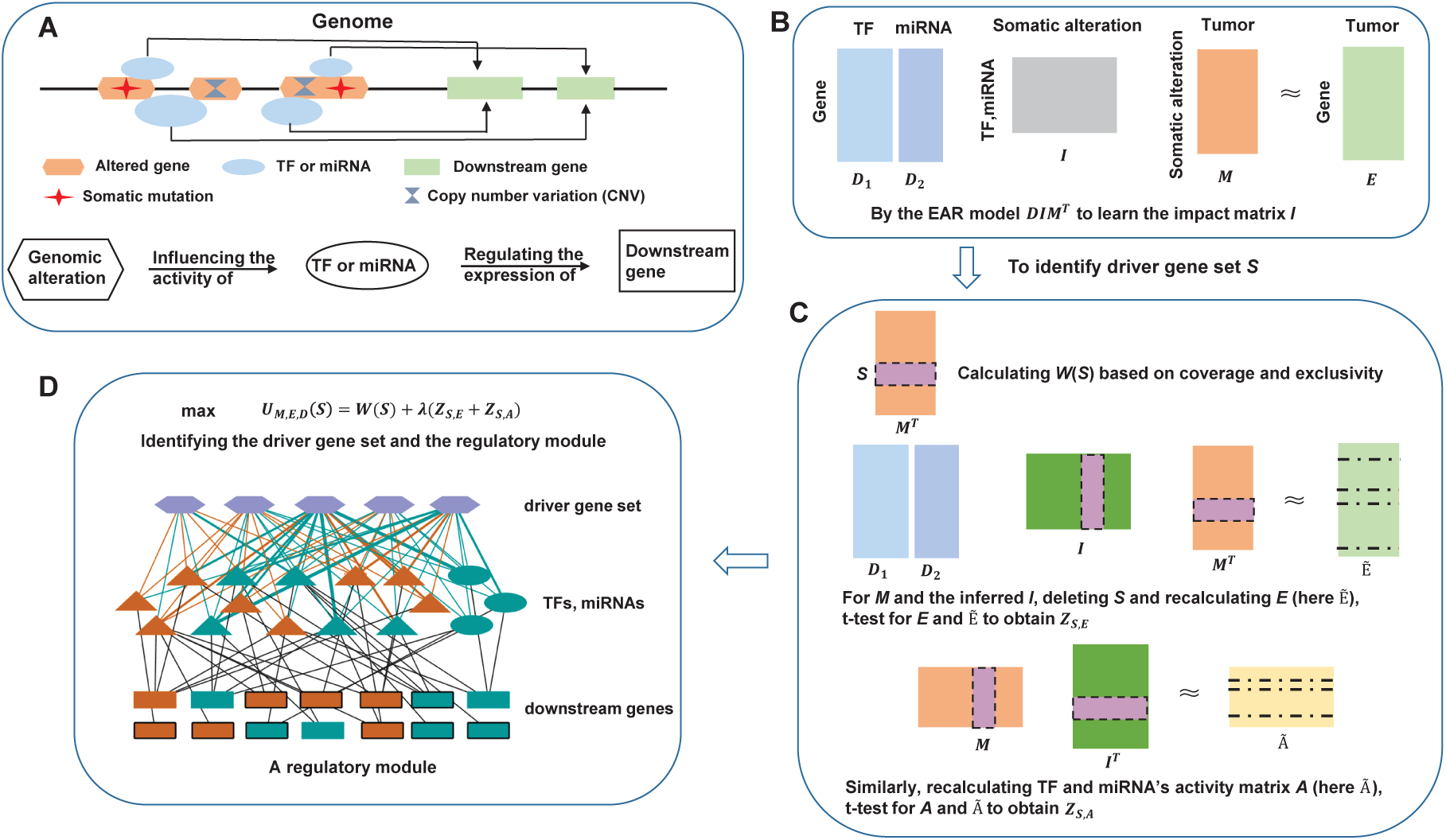
Overview of the main idea and workflow in this study. A) A schematic diagram for gene regulation; B) The integrated data (including genomic, transcriptomic and epigenetic data) and the characterization of their relationships; C) Calculating the impacts of an altered gene set on activities of TFs and miRNAs, respectively; D) Establishing an optimized model to identify the driver gene set and the regulatory module (a three-layer network).

For a gene set *S* in the mutation matrix *M*, keeping in mind that *M* and *E* have the relation in Eq. 2, by deleting *S* from *M* and recalculating Eq. 2 we obtain a new gene expression matrix *Ẽ*. Performing *t*-test on *E* and *Ẽ*, we denote *P_S,E_* as the set of all *p*-values less than a certain threshold *β* (for example, *β*=0.001), which corresponds to differentially expressed genes. Suppose *P_S,E_* = {*p_S,E_*(1)*, p_S,E_*(2)*, . . ., p_S,E_*(*k*_1_)} with the set size *k*_1_. Then the Stouffer’s *Z*-score [18], which is closely related to Fisher’s combined *p*-value [19] and reflects a overall extent of significance of the *P_S,E_*, can be got as follows:

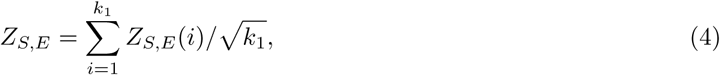

where *Z_S,E_*(*i*) = Φ*^−^*^1^(1 *− p_S,E_*(*i*)), and Φ is the standard normal cumulative distribution function.

Similarly, deleting *S* from *M* we recalculate Eq. 3 and obtain a new activity matrix *Ã*. Performing *t*-test on *A* and *Ã*, we denote *Q_S,A_* as the set of all *p*-values less than *β*, which corresponds to TFs and miRNAs with significantly changed activities, and set *Q_S,A_*= *{q_S,A_*(1)*, q_S,A_*(2)*, . . ., q_S,A_*(*k*_2_)*}* with the set size *k*_2_. Then we get the Stouffer’s *Z*-score [18] for TFs and miRNAs:

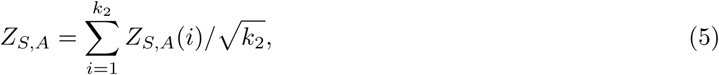

where *Z_S,_* (*i*) = Φ*^−^*^1^(1 *− q_S,A_*(*i*)).

We introduce **gtePIDP** (genomic-transcriptomic-epigenetic Phenotypically Impactful Driver Pathways), a computational framework that integrates somatic mutations, gene expression, and TF/miRNA regulatory networks to identify driver pathway or gene set (here, *S*) as follows:

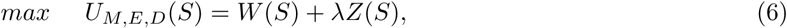

where *M*, *E*, *D*, and *W* (*S*) are stated as above; *Z*(*S*) = *Z_S,E_* + *Z_S,A_* in which *Z_S,E_*and *Z_S,A_* are defined in Eqs. 4 and 5, respectively; and *λ* is a parameter trading off between gene mutations and their impacts on phenotypes in identifying driver pathways in cancer. We investigated the influence of the parameters (*λ* and *β*) on the performance of the model gtePIDP (Eq. 6) in the **Supplementary information** (Tables S1 S12). We solve gtePIDP (Eq. 6) by Genetic Algorithm (The code for the Matlab package of gtePIDP is freely available at https://github.com/zjh136/project-gtePIDP).

### 2.4 Biological data

The mutation dataset and corresponding RNA-seq expression profiles were obtained from TCGA program through distinct acquisition pipelines. The curated mutation data were directly acquired from a previous comprehensive study [20], and the RNA-seq data (Illumina HiSeq 2000 platform) were retrieved via the Genomic Data Commons (GDC) portal (https://portal.gdc.cancer.gov/). TF-gene regulatory interactions and miRNA-target relationships were downloaded from the Molecular Signatures Database (https://www.gsea-msigdb.org/gsea/msigdb).

## 3 Results

In view of the complex molecular heterogeneity and poor prognosis of GBM, here we use the proposed model gtePIDP (Eq. 6) to deeply investigate the molecular regulatory mechanism hidden in the heterogeneous alterations and the phenotypes of the tumors, which may provide some useful information for the effective treatment of this cancer. Within this process we also analyze the influence of the parameters on the performance of the model. Furthermore, we validate the effectiveness of gtePIDP by randomly selecting gene sets as the candidate ones.

### 3.1 The gtePIDP can effectively identify functional driver gene set and the related regulatory module with TFs and miRNAs

In gtePIDP (Eq. 6) the parameter *λ* directly trades off the weight *W* and the *Z*-score *Z_S,E_* + *Z_S,A_*, and the parameter *β* is the threshold of *p*-values. In the **Supplementary information** we investigated the influence of these parameters on the performance of the model by respectively choosing *λ* = 0.1, 0.3, 0.5 and *β* = 0.0001, 0.001, 0.005, 0.01 for GBM (Tables S1 S12).

From these results we notice that both *λ* and *β* trade off somatic mutations and their impacts on phenotypes (*W* and *λ · Z*, respectively) in identifying driver pathways. Along with *λ* or *β* getting larger we can detect more genes and more TFs and miRNAs, but the identified driver gene sets are similar (for example, **Supplementary information**, Tables S1, S2).

In order to get the driver gene sets with large exclusivity and coverage (i.e., large *W*), and at the same time discover more and significant genes, TFs and miRNAs (with respect to differential expressions and changed activities, respectively), we use *β* = 0.0001*, λ* = 0.3 for the following analysis (Table 1).

**Table 1.**
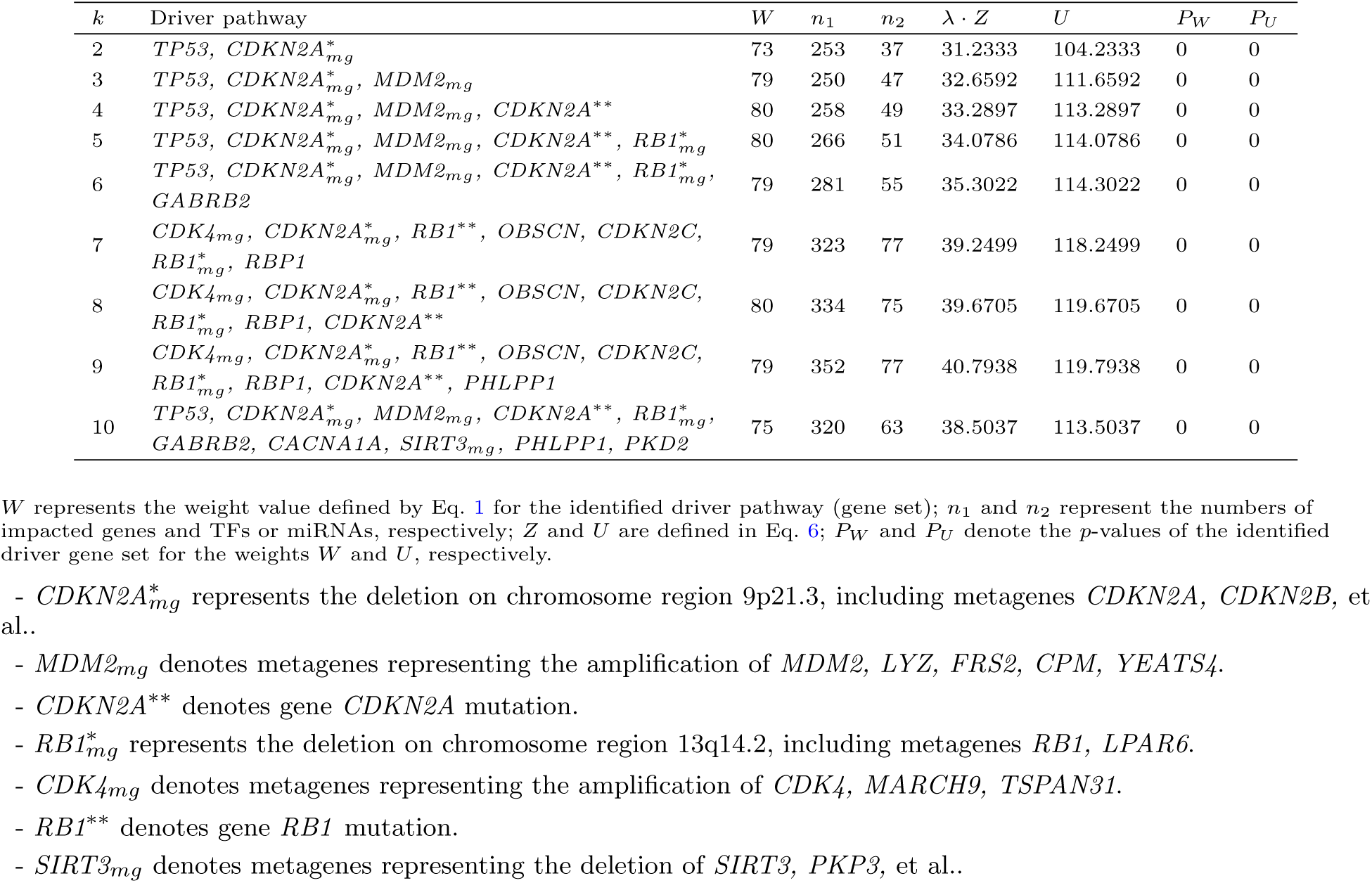
Identified significant driver gene sets for *β* = 0.0001 and *λ* = 0.3 in GBM.

From Table 1 we see that using gtePIDP we not only identify gene sets involved in important biological signalling pathways, more importantly, we detect TFs and miRNAs which in turn regulate the downstream gene expressions. This is very different from many previous studies for discovery of driver gene sets, such as Dendrix [3], MDPFinder [21] and Multi-Dendrix [22], which only identify candidate driver gene sets, but do not consider their impacts on the activities of TFs and miRNAs, as well as on the expressions of downstream genes.

Furthermore, as a comparative study, we wonder if gtePIDP can also find biologically meaningful results on randomly selected gene sets. To this end, we investigate the performance of gtePIDP on randomly selected gene sets in GBM. Without loss of generality, we take *k* = 5 as an example for the analysis, and the results of 10 runs are displayed in Table S13 (**Supplementary information**). Note that for this study we simultaneously consider two kinds of significance, i.e., *P_W_* -value and *P_U_* -value. For all the 10 runs in Table S13, *P_W_* -value is larger than 0.05, so all the detected gene sets cannot be candidate driver gene sets although for the ninth run the *P_U_* -value is less than 0.05. This comparative study validates the feasibility and effectiveness of the proposed method for identifying driver pathways and exploring possible pathogenesis in cancer (Table 1).

In the following, without loss of generality, for GBM we will take *k* = 5 as an example to demonstrate the functions of the identified genes, TFs and miRNAs, as well as their relevance to carcinogenesis.

For *k* = 5, we identify the driver pathway including genes *TP53,* 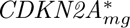*, MDM2_mg_, CDKN2A^∗∗^,* 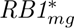, which are involved in the DNA damage response and cellular apoptosis pathway (Table 1, Fig. 2A). In addition, we detect 51 TFs and miRNAs with significantly changed activities and 266 significantly differentially expressed genes (Fig. 2B).

**Fig. 2.**
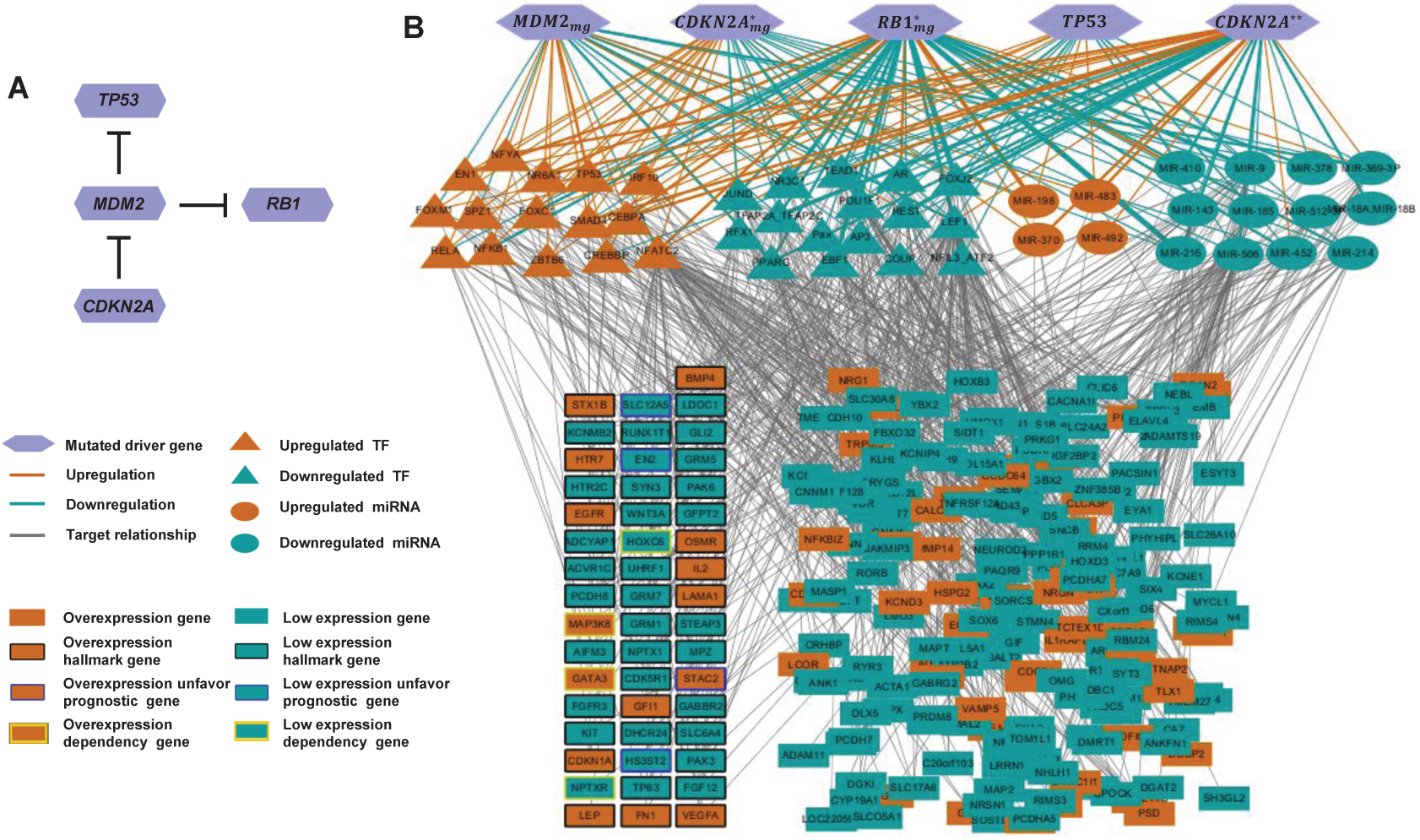
A) The driver pathway identified by gtePIDP for *k* = 5 in GBM. B) The identified driver gene set and its regulation for the activities of TFs, miRNAs, and the expressions of the downstream genes in GBM. The thickness of the brown and cyan lines reflects the strength of the impacts of the mutated genes on TFs and miRNAs, according to the scales on the values of *I^∗^* in Eq. 7.

Hereafter in this study, for each detected TF or miRNA (regulator), if the number of samples with increased activities larger than that with decreased activities, we say the TF or miRNA has up-activity, otherwise we say it has down-activity (Fig. 2B, Fig. 3A). Similarly, for each detected downstream gene, if the number of samples with over-expression is larger than that with low-expression, we say it is upregulated, otherwise we say it is downregulated. In fact, here for TF or miRNA, these up-activities or down-activities are mainly determined by the impacts of the alterations of the driver genes on these regulators. For example, most of the weights of impacts of the driver genes on *FOXJ2* are negative, resulting in *FOXJ2* down-activity; whereas most of the weights of impacts of the driver genes on *NFATC2* are positive, resulting in *NFATC2* up-activity (some of the relations are marked by dashed lines in Fig. 3A,B). Furthermore, to catch hold of the important relations and thus get rid of weak links between the mutated driver genes and regulators, we introduce a max-min threshold to make the impact matrix sparser. Specially, for *k* identified driver genes, suppose we find *l* TFs and/or miRNAs with significantly changed activities, then we obtain an *l × k* impact matrix *I̅*. Let *Ĩ* denote the absolute value of *I̅*. For each TF or miRNA *i*, set *V_i_* = *max_j_Ĩ*(*i, j*) and *V* = *min V_i_*. For the max-min threshold *V*, we define the sparser matrix *I^∗^* as follows (Fig. 3C)

**Fig. 3.**
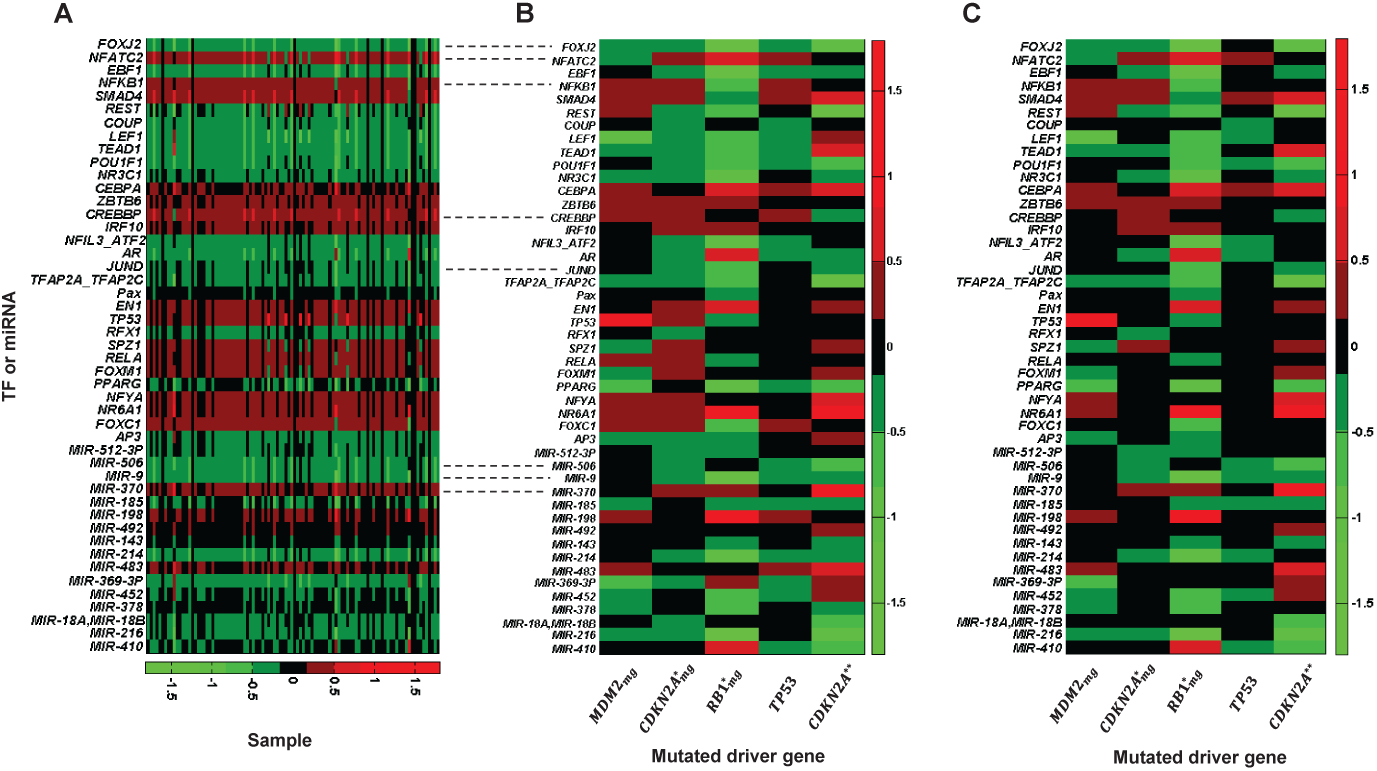
A) TFs and miRNAs with significantly changed activities in GBM. B) The weights of the impact of the driver pathway on the TFs or miRNAs in GBM. C) The sparser weights defined by Eq. 7 which may correspond to the important impacts of the driver pathway on the TFs or miRNAs in GBM.

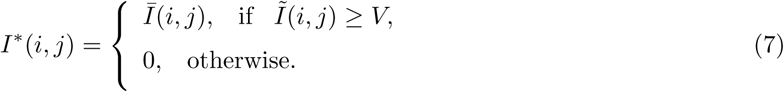

Furthermore, to differentiate the strength of the impacts of driver genes on TFs or miRNAs, we scale the absolute values of *I^∗^* by four intervals: [V, 0.5), [0.5, 1.0), [1.0, 1.5), and larger than 1.5 (Fig. 2B, Fig. 3C).

### 3.2 The TFs and miRNAs with significantly changed activities are relevant to carcinogenesis

Accumulating research indicates that TFs and miRNAs play critical roles in carcinogenesis of different tumors and other complex diseases. The present study is designed to identify cancer candidate driver gene sets and further investigate their impacts on TFs and miRNAs as well as the downstream gene expressions (phenotypes), with an expectation to provide proper clues on the regulatory mechanisms and therapeutic applications.

In Fig. 2 there are 31 TFs and 16 miRNAs, each of which with up-activity or down-activity (Fig. 3A). Many of them are important regulators, and are closely related to cancer development and progression. For example, we infer *NFATC2* and *MIR-506* have up-activity and down-activity in GBM, respectively, and they affect a lot of downstream differentially expressed genes (71/266 and 43/266, respectively) (Fig. 4A,B). *NFATC2* is a member of the nuclear factor of activated T cells (NFAT) family, which plays a central role in inducing gene transcription during the immune response. A lot of studies indicated that *NFATC2* and *MIR-506* are highly involved in the carcinogenesis and progression of various cancers [23–25], including glioblastoma, breast cancer, and so on. For instance, Zuo *et al.* demonstrated that *NFATC2* expression was significantly increased in both GBM tissues and cultured cell lines, and *NFATC2* knockdown in GBM cells obviously inhibited cell proliferation and elevated apoptosis [26]; Luo *et al.* revealed that *MIR-506* inhibited the proliferation and invasion by targeting *IGF2BP1* in GBM [27]; and Peng *et al.*showed that over-expression of *MIR-506* markedly suppressed cell proliferation, colony formation, migration and invasion in glioma cells [28]. Taken together, all these showed that *NFATC2* is an oncogene and *MIR-506* is a tumor suppressor, which is quite consistent with the discovery in this study, where we find *NFATC2* and *MIR-506* are upregulated and downregulated in tumors (Fig. 3A,B), respectively.

**Fig. 4.**
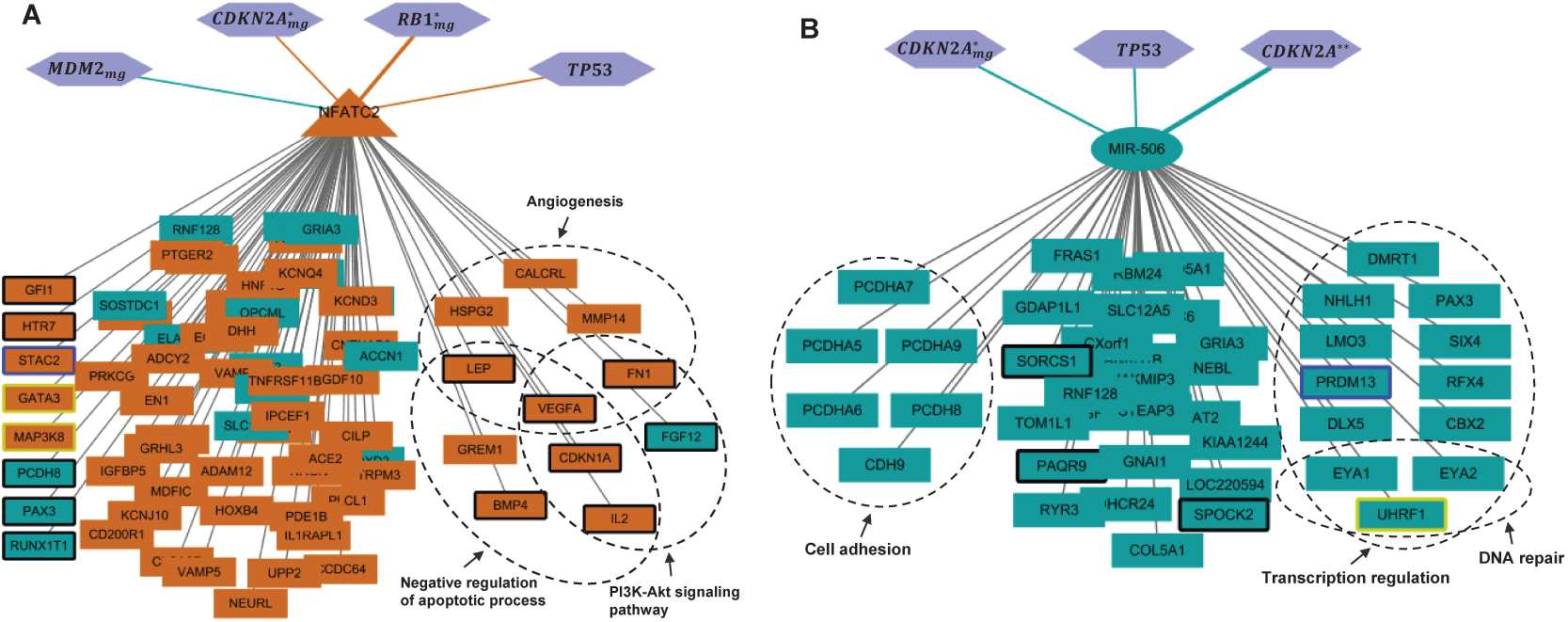
The regulators and their affected genes in GBM: A) *NFATC2*, B) *MIR-506*.

Besides the regulators with large numbers of targets, whereas some other regulators have medium or few target genes, for example, *FOXJ2, MIR-9* and *MIR-370* affect 15, 26 and 4 differentially expressed genes, respectively. All of them are also associated with carcinogenesis. *FOXJ2* has been found to be involved in tumor migration and invasion as well as epithelial-mesenchymal transition (EMT) in various cancers. Low *FOXJ2* expression was observed in human glioma tissues and the expression level was correlated with the grade of malignancy, and over-expression of *FOXJ2* significantly inhibited glioma cell migration [29]. Many studies in cancer cells demonstrated that *MIR-9* was a tumor suppressor in multiple cancers including GBM by targeting various mRNAs [30–32]. Specially, *MIR-9* induced decreased ability of GBM cells to migrate and invade via regulation of *MAPK14* signaling elements [30]. *MIR-9* over-expression triggered the apoptosis of GBM cell lines via suppressing *SMC1A* expression [31]. In this study, the activities of the tumor suppressors *FOXJ2* and *MIR-9* are downregulated in GBM (Fig. 3A,B). All these findings strongly suggest that *FOXJ2* and *MIR-9* are candidate prognostic markers and interesting therapeutic targets for GBM, especially *MIR-9* suppresses cancer cell growth through multiple pathways.

As for *MIR-370*, a great deal of evidence has shown that *MIR-370* plays an important role in the development and progression of tumor [33, 34]. Here this study indicates that *MIR-370* may act as an onco-miRNA in GBM, where the activity is upregulated for tumorigenesis (Fig. 3A,B).

### 3.3 The downstream affected genes are involved in cancer biological processes, for which we find their regulatory modules

First, some of the 266 downstream affected genes are cancer hallmark genes, prognostic genes or cancer dependent genes.

#### Hallmark genes

The hallmarks of cancer has provided an important molecular framework for deeper understanding cancer pathogenesis, and from the corresponding cellular processes, 2172 genes have been defined as hallmark-related genes [35]. We find that there are 43 genes are hallmark-related in the downstream affected genes (the genes with black boxes in Fig. 2B).

#### Prognostic genes

Uhlen *et al.* used a systems level approach to analyze the transcriptome of 17 major cancer types with respect to clinical outcome, and for each cancer type they identified candidate prognostic genes associated with clinical outcome [35]. In the detected genes there are five genes showing unfavorable prognostic effects (that is, high expression results in short survival of the patients) (the genes with blue boxes in Fig. 2B).

#### Cancer dependent genes

Tsherniak *et al.* performed systematic loss of function screens in a large number of well-annotated cell lines, and they identified candidate genes essential for cancer cell proliferation/survival (cancer dependent genes) [36]. There are five cancer dependent genes in the downstream affected genes (the genes with yellow boxes in Fig. 2B).

Then we use DAVID [37] to annotate functions of the 266 downstream affected genes in GBM. Some of the genes are involved in important biological processes or signaling pathways related to cancer hallmarks. Especially, we notice that the important regulators *NFATC2* and *MIR-506* regulate the downstream genes in several processes or pathways, such as angiogenesis and DNA repair, respectively (Fig. 4). However, these processes or pathways may be regulated by many regulators simultaneously. Here we can use the proposed method to identify the regulators, and more importantly, to infer the influence extent of the alterations of the mutated genes to the activity of the regulators.

Sustained angiogenesis is one of the important cancer hallmarks [38]. For the six genes (*CALCRL, FN1, HSPG2, LEP, MMP14, VEGFA*) related to angiogenesis (Fig. 4A) we identify the regulatory module in Fig. 5.

**Fig. 5.**
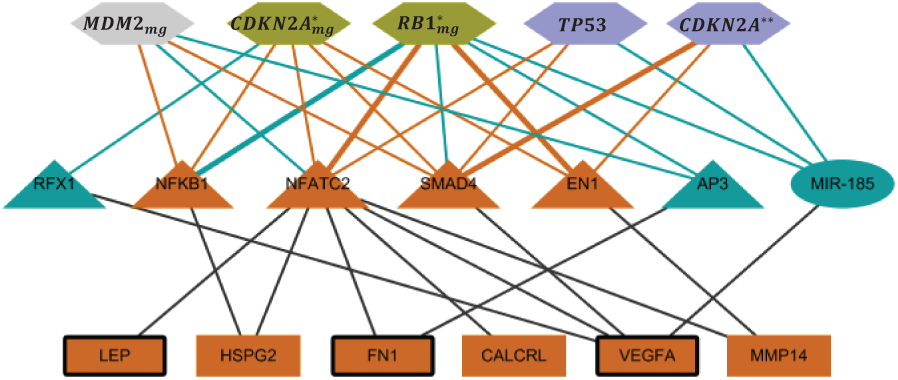
The regulatory module related to angiogenesis in GBM.

Besides *NFATC2* stated in last subsection, other regulators may also play important roles in the process of angiogenesis. *NFKB1* is involved in a series of signal transduction events related to many biological processes, and its activation is an important driver of the malignant phenotype that confers a negative prognosis in patients with GBM [39]. *EN1* participates in the development of the brain. High expressions of *EN1* exist in various tumors, especially, its overexpression may be a potential prognostic marker in glioma [40]. *SMAD4* is involved in the TGF-*β* signaling, which has a dual function in carcinogenesis, i.e., either promotes or inhibits tumor formation and progression of many cancer types [41]. Here we speculate that *SMAD4* has the role of promoting progression of GBM based on its upregulation resulting in the overexpression of the hallmark gene *VEGFA* (Fig. 5). Later we found that this inference is consistent with the recent study [42], where TGF-*β*/SMAD4 was shown to be involved in glioblastoma migration and invasion. *AP3*, as an artificially designed peptide, exhibited potent antiviral activity against a broad spectrum of HIV-1 strains [43]. *RFX1* has been shown to be a tumor suppressive transcription factor in GBM, and its upregulation can prevent the proliferation of glioblastoma cells [44]. At last, a lot of studies revealed that *miRNA-185* inhibits proliferation, migration and invasion in various types of cancer [45, 46]. Moreover, Ma *et al.* demonstrated that *miRNA-185* inhibits cell proliferation and induces cell apoptosis by targeting *VEGFA* in the clear cell renal cell carcinoma [47]. All these studies indicate that *NFATC2, NFKB1, EN1* and *SMAD4* are oncogenes, and *AP3, RFX1* and *MIR-185* are tumor supressors. These conclusions are quite consistent with our study (Fig. 5), where the activities of the first four TFs are upregulated and those of the last three TFs or miRNAs are downregulated in GBM tumors. We suggest that *MIR-185/VEGFA* might be a valuable therapeutic target for GBM.

### 3.4 Through the three-layer network we can infer the candidate therapeutic axes

By gtePIDP we can construct a three-layer network for the cancer (Fig. 2B). The above analysis indicates that the identified genes and miRNAs are highly involved in carcinogenesis of the cancer (here GBM). In fact, anti-angiogenic therapies are an important class of anti-cancer treatment. It has exhibited excellent efficacy against some human tumours, such as colorectal cancer and non-small cell lung cancer. GBM showed high extent of neovascularization, but the existed performance of anti-angiogenic therapies is not satisfactory [48, 49]. So it is vitally necessary to identify new molecular biomarkers to be considered as new targets. The above mentioned discovery may provide some useful information for the target treatments of GBM. Moreover, by the three-layer network we can also provide some candidate therapeutic axes for GBM.

#### *TP53/MIR-185/VEGFA* axis

In general, targeting perturbed tumour pathways is the key idea in individualised cancer therapy. It is well-known that *TP53* is a tumor suppressor gene frequently mutated in cancer and implicated in cell-cycle regulation, apoptosis, and angiogenesis. Joshi *et al.* studied the pathways disrupted in association with *TP53* mutation status in breast cancer, and they identified *VEGFA* expression as an important marker of survival for the patients [50]. Schwaederle *et al.* demonstrated that *TP53* mutations are associated with higher *VEGFA* expression for patients with non-small cell lung cancer, and investigated the improved outcomes after anti-angiogenic treatment for patients with mutant *TP53* versus wild-type *TP53* [51]. Recently, Li *et al.* further studied the relationship between *TP53* -mutation status and *VEGFA* expression by pan-cancer analysis (30 cancer types, including GBM) [52]. They found statistically significant increases of *VEGFA* expression in *TP53* -mutated adenocarcinomas compared to *TP53* wild-type tumors, which provided additional evidence that *TP53* mutations are linked to the *VEGF* pathway. Based on the identified functional module with angiogenesis for GBM (Fig. 5), in view of *MIR-185* inhibiting cell proliferation and inducing cell apoptosis by targeting *VEGFA* in some cancer stated above [47], we infer that *TP53/MIR-185/VEGFA* may be a candidate anti-angiogenic therapeutic axis for GBM, which deserves in depth further studies from therapeutic and prognostic context.

#### *MDM2/SMAD4/VEGFA* axis

*MDM2* is a key inhibitor of *TP53* and a positive activator of *VEGF* activity with an important role in neuroblastoma pathogenesis [53]. Furthermore, the suppression of *VEGF* expression subsequent to inhibition of *MDM2* in *TP53* mutant cells suggests that *MDM2* has a regulatory role on *VEGF* expression through a *TP53* -independent mechanism [54]. In addition, a previous study indicated that *SMAD4* participated in activating the expression of *VEGFA* and was related to the progression of colorectal cancer [55]. Moreover, by the proposed method we observed that the amplification of *MDM2* could increase the activity of *SMAD4* (Fig. 5). Together, these indicate that *MDM2/SMAD4/VEGFA* may be another candidate anti-angiogenic therapeutic axis for GBM.

### 3.5 Candidate therapeutic axes related to PI3K-Akt signaling in GBM

Previous studies indicate that among others, the *TP53* and PI3K-Akt pathways are all involved in GBM [13]. Here we can use the proposed model gtePIDP to identify a module which shows the links how the driver *TP53* pathway alterations affect the expression of genes in the PI3K-Akt pathway (Fig. 6). Moreover, through this model we can also find some candidate pathogenetic axes for cancers.

**Fig. 6.**
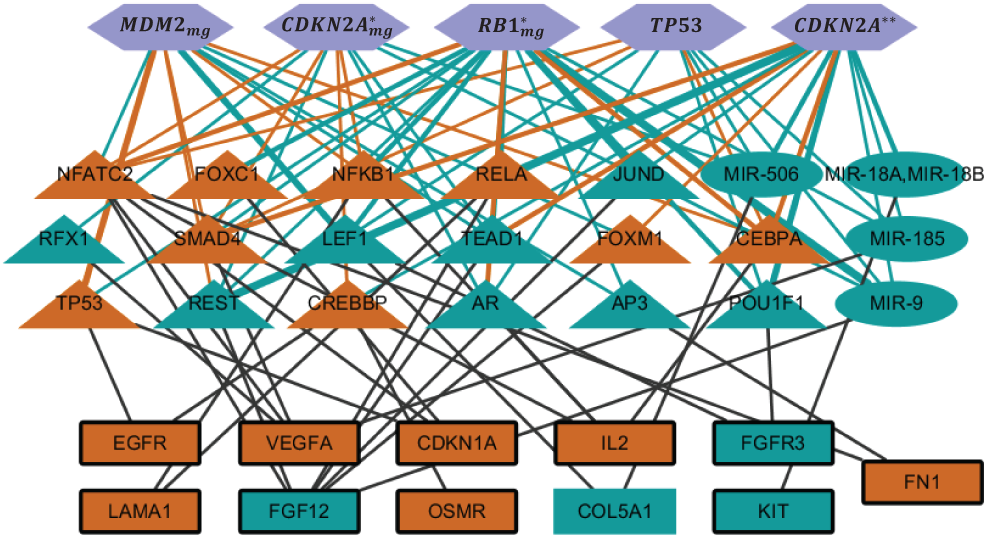
The regulatory module related to PI3K-Akt signaling in GBM.

#### *CDKN2A/CREBBP/OSMR(/EGFR)* axis

*OSMR* is a cytokine receptor gene of IL-6 family and has been reported to play an important role in driving the glioma cells towards mesenchymal type, so it can be as a predictive factor for progressive malignancy in GBM [15], and its overexpression is correlated with poor survival for GBM patients [58]. *OSMR* is also shown to be a prognostic marker in the response prediction to radiotherapy and chemotherapy and contributes to the regulation of local immune response in GBM [59]. Furthermore, previous studies indicate that *CREBBP* is involved in promoting cell proliferation and anti-apoptosis in a lot of cancers, including GBM and Glioma [60, 61]. Sung *et al.* demonstrated that *CDKN2A* is usually deleted in primary central nervous system lymphoma (PCNSL), and *OSMR* is significantly differentially expressed between PCNSL and normal lymph node tissues [62]. Here in this study in the identified functional module (Fig. 6), *OSMR* and *EGFR* are just two upregulated target genes of *CREBBP*, which has the up-activity by the deletion and mutation of *CDKN2A*. Taken together, we suggest that *CDKN2A/CREBBP/OSMR(/EGFR)* might be a promising therapeutic axis for GBM.

#### *TP53(/CDKN2A/RB1)/MIR-9/FGF12* axis

*FGF12* encodes a member of the fibroblast growth factor (FGF) family, which are involved in a variety of biological processes, including tumor growth and invasion. Although it was overexpressed in some cancers (such as esophageal squamous cell carcinoma [63]), there are several other kinds of cancers in which *FGF12* was downregulated or hypermethylated, such as in breast cancer [64]. As stated above, *MIR-9*, as a tumor suppressor, played critical roles for tumorigenesis with its downregulated activity in GBM [30, 31]. Here in the identified functional module (Fig. 6), *FGF12* as the only target of *MIR-9*, its expression is also downregulated, indicating *FGF12* is also a tumor suppressor in GBM. On the other hand, the genomic mutations or deletions of *TP53, CDKN2A* and *RB1* induced the down-activity of *MIR-9* (Fig. 6), and there are studies demonstrated that these genes may cooperate with *MIR-9* to participate the biological process of tumorigenesis [65–67]. All these may indicate that *TP53(/CDKN2A/RB1)/MIR-9/FGF12* might be another promising therapeutic axis for GBM.

Other more complicated regulatory modules can also be identified and constructed by gtePIDP, such as the functional module related to apoptosis in GBM (**Supplementary information**, Fig. S1). Taken together, we can use the proposed model to identify candidate driver gene sets, to investigate the possible pathogenesis by considering their impacts on the activities of TFs and miRNAs, and further explore their influence on phenotypes, which may provide possible candidate drug targets for cancer therapy.

## 4 Discussion

The molecular complexity of GBM poses significant challenges in pinpointing driver pathways that underlie tumor progression and therapeutic resistance. Traditional computational methods, such as MutSig and Dendrix, prioritize genes with high mutation rates or mutual exclusivity but often overlook the downstream regulatory cascades that mediate phenotypic impact. Here, we present gtePIDP (Eq. 6), a novel framework that integrates genomic, transcriptomic, and epigenetic data to identify driver pathways with both mutational coherence and functional consequences. Our approach not only refines driver gene set prediction but also unravels mechanistic links between somatic alterations and transcriptional/post-transcriptional dysregulation, offering a paradigm shift in cancer genomics.

A key strength of gtePIDP lies in its dual evaluation of mutation patterns (*W* (*S*)) and phenotypic perturbations (*Z*(*S*)). By incorporating regulatory network analysis via an extended affinity regression (EAR, Eq. 2) model, we quantified how mutations in driver gene set (*TP53, CDKN2A, MDM2, RB1*) propagate to disrupt TF and miRNA activities, ultimately rewiring oncogenic pathways. For instance, *TP53* alterations were found to suppress *MIR-185*, leading to *VEGFA* overexpression – a critical mediator of angiogenesis. This axis aligns with pan-cancer evidence linking *TP53* mutations to elevated *VEGFA* expression and poor prognosis [52], yet our study uniquely identifies *MIR-185* as a novel intermediary in GBM. Similarly, *MDM2* amplification upregulated *SMAD4* activity, further activating *VEGFA* and highlighting a *TP53* -independent angiogenic mechanism. These findings underscore the value of integrating regulatory networks to uncover context-specific vulnerabilities.

The identified PI3K-Akt signaling modules, including *CDKN2A/CREBBP/OSMR(/EGFR)* and *TP53(/CDKN2A/RB1)/MIR-9/FGF12*, provide insights into GBM’s mesenchymal transition and therapy resistance. *OSMR*, a cytokine receptor linked to immune evasion and poor survival [58, 59], emerged as a downstream effector of *CREBBP* activation driven by *CDKN2A* deletions. This aligns with prior work implicating *OSMR* in promoting mesenchymal phenotypes and radioresistance [15, 59]. Meanwhile, the tumor-suppressive *MIR-9*, silenced by *TP53* and *RB1* alterations, was shown to repress *FGF12* – a potential PI3K-Akt inhibitor. These results resonate with studies demonstrating *MIR-9* ’s role in suppressing glioma migration and apoptosis [30, 31], yet our framework uniquely maps its regulation to upstream genetic lesions, offering actionable targets for combinatorial therapies.

Our findings also emphasize miRNA dysregulation as a hallmark of GBM pathogenesis. For example, *MIR-506* downregulation, driven by driver mutations, correlated with oncogenic pathway activation, consistent with its reported tumor-suppressive roles in glioma [27, 28]. Conversely, *MIR-370* upregulation promoted proliferation, mirroring its oncogenic functions in other cancers [33, 34]. Such regulatory hierarchies, inferred via gtePIDP, highlight miRNAs as pivotal nodes connecting genetic alterations to phenotypic outcomes.

In addition to GBM, the gtePIDP framework is readily extendable to other malignancies such as BRCA, OV, LUAD, and so on (**Supplementary information**), providing a universal computational strategy for pan-cancer biomarker discovery. For instance, the identified driver pathways are listed in Tables S14 and S15 for BRCA and OV, respectively. Take *k* = 8 as an example. For BRCA, we identified *TP53, PIK3CA, GATA3, CDH1, CTCF, FLRT3mg, MYCNmg, MAPK1* as the driver gene set, which influenced activity changes of 32 TFs or miRNAs, and differential expressions of 176 downstream genes. While for OV, the disturbed TFs and/or miRNAs as well as downstream genes are much more than those for BRCA. Here we identified *TP53, ATP1A1OSmg, KRAS, NRAS, FAM124B, CDK17, FLT3, MAP3K1* as the driver gene set, which influenced activity changes of 105 TFs or miRNAs, and differential expressions of 556 downstream genes. We notice that there are a large proportion of disturbed genes encoding glycoproteins in these two types of cancers (42.61% and 43.58%, respectively), which indicates that glycoproteins may play crucial roles in these cancers. Another important factor is alternative splicing, to which 45.45% and 46.11% of genes are related, respectively. In addition, we also find that some of the disturbed genes in these two types of cancers are related to cell-cell adhesion, and some are involved in the MAPK signaling pathway, transmembrane receptor protein tyrosine kinase signaling pathway and pathways in cancer. All these demonstrate that BRCA and OV indeed have some commonalities in carcinogenesis. But on the other hand, here we also find some difference between them. For example, response to oxidative stress is observed in BRCA other than OV; whereas Wnt signaling pathway, positive regulation of protein amino acid phosphorylation, Hippo signaling pathway, cell migration, angiogenesis and Rap1 signaling pathway are observed in OV other than BRCA. This indicates that although previous studies manifested that there are some commonalities between BRCA and OV, much difference exists for the impact of driver genes on TFs and miRNAs as well as their regulation on downstream gene expressions. From this point of view we may suggest that OV might have much more complex pathogenesis than BRCA.

Despite these advances, certain limitations warrant consideration. First, gtePIDP relies on predefined TF-miRNA-gene networks, which may omit context-specific interactions. While we employed stringent thresholds (e.g., max-min sparsification) to prioritize robust connections, future iterations could benefit from dynamic network modeling. Second, parameter selection (e.g., *λ, β*) influences pathway prioritization, necessitating careful optimization across datasets. Third, while functional enrichment of downstream genes (e.g., angiogenesis, apoptosis) supports biological relevance, experimental validation of predicted axes (*TP53/MIR-185/VEGFA, MDM2/SMAD4/VEGFA*) is critical to confirm causality.

Future directions should focus on translating these computational insights into preclinical models. For instance, targeting *VEGFA* via *MIR-185* mimics or *SMAD4* inhibitors could test the efficacy of antiangiogenic strategies in *TP53* -mutant or *MDM2* -amplified GBM subtypes. Similarly, restoring *MIR-9* expression in therapy-resistant tumors may sensitize cells to PI3K-Akt inhibitors. Experimental validations will bridge the gap between computational discovery and clinical implementation.

## 5 Conclusion

In this study, we developed gtePIDP, a computational framework that integrates multi-omics data to identify driver pathways with maximal phenotypic impact in cancers. By bridging somatic mutations with transcriptional and post-transcriptional regulatory consequences, gtePIDP uncovered key driver gene set for GBM (*TP53, CDKN2A, MDM2, RB1*) and its dysregulated networks involving oncogenic (*NFATC2, MIR-370*) and tumor-suppressive regulators (*MIR-506, MIR-9, FOXJ2*). Involved pathways drive critical hallmarks of GBM, including angiogenesis, apoptosis evasion, and proliferation, while novel therapeutic axes (*TP53/MIR185/VEGFA, MDM2/SMAD4/VEGFA, CDKN2A/CREBBP/OSMR(/EGFR)*) offer mechanistic insights into angiogenic signaling, mesenchymal transition and therapy resistance.

Our findings underscore the importance of regulatory network analysis in deciphering cancer complexity. The suppression of *MIR-185* and *MIR-9* by *TP53* and *RB1* alterations, respectively, highlights miRNA dysregulation as a central feature of GBM pathogenesis. Similarly, the role of *OSMR* and *CREBBP* in mediating therapy resistance aligns with prior clinical observations, validating gtePIDP’s predictive power.

While the study advances driver pathway identification, experimental validation of predicted targets and further refinement of regulatory networks remain essential next steps. Translating these insights into targeted therapies – such as *MIR-185* replacement or *SMAD4* inhibition – could address unmet needs in GBM treatment.

In conclusion, because gtePIDP is readily extendable to other malignancies (such as BRCA, OV, LUAD, etc.), it provides a robust, systems-level approach to unravel the molecular underpinnings of cancers. By prioritizing pathways with both mutational and functional significance, this framework not only enhances our understanding of tumorigenesis but also may pave a way for tailored therapeutic strategies in these lethal diseases.

## Data availability

All biological data used in this study are available on public data platforms. The GBM mutation dataset and corresponding RNA-seq expression profiles were obtained from TCGA program ([20], https://portal.gdc.cancer.gov/). TF-gene regulatory interactions and miRNA-target relationships were downloaded from the Molecular Signatures Database (https://www.gsea-msigdb.org/gsea/msigdb).

The code for the Matlab package of gtePIDP is freely available at https://github.com/zjh136/project-gtePIDP.

## Acknowledgements

We would like to thank Professor Shihua Zhang from the Academy of Mathematics and Systems Science, CAS for the suggestion.

