## Supplemental Tables S1-S16, Supplemental Figure S1 for "Integrative multi-omics framework identifies phenotypically impactful driver pathways in glioblastoma multiforme"

Junhua Zhang

Academy of Mathematics and Systems Science, Chinese Academy of Sciences, Beijing 100190, China.

In this study we developed an integrated model gtePIDP to identify cancer driver pathways where somatic alterations, gene expressions, transcription factor (TF) regulatory networks as well as microRNA (miRNA) targeting relations are combined.

#### 1 Investigation on the selection of $\lambda$ and $\beta$

In the proposed model the parameter $\lambda$ directly trades off the weight $W$ (involving exclusivity and coverage of mutations) and the $Z$-score $Z_{S,E} + Z_{S,A}$ (reflecting the impact of gene mutations on TFs and miRNAs as well as downstream gene expressions), and the parameter $\beta$ is the threshold of $p$-values, where only those less than $\beta$ are considered for the calculation of the $Z$-score. Here taking glioblastoma multiforme (GBM) data as an example we investigate the influence of these parameters on the performance of the model by choosing $\lambda = 0.1, 0.3, 0.5$ and $\beta = 0.0001, 0.001, 0.005, 0.01$, respectively (Tables S1 - S12).

From these results we notice that both $\lambda$ and $\beta$ trade off somatic mutations and their impacts on phenotypes ($W$ and $\lambda \cdot Z$, respectively) in identifying driver pathways. Along with $\lambda$ or $\beta$ getting larger we can detect more genes (with differential expressions) and more TFs and miRNAs (with significantly changed activities), but the identified driver gene sets are similar. For example, for $\beta = 0.0001, \lambda = 0.1$ (Table S1) and $\beta = 0.001, \lambda = 0.1$ (Table S2) we identify exactly the same driver gene sets, the only difference is the numbers of detected genes, TFs and miRNAs. In fact, $\lambda$ and $\beta$ have similar performance on the gtePIDP model for other types of cancers, such as breast carcinoma (BRCA), ovarian cancer (OV), lung adenocarcinoma (LUAD), etc.

In order to get the driver gene sets with large exclusivity and coverage (i.e., large $W$), and at the same time discover more and significant genes, TFs and miRNAs (with respect to differential expressions and changed activities, respectively), in this study we use $\beta = 0.0001, \lambda = 0.3$ for the following analysis.

**Table S1** Identified significant driver gene sets for $\beta = 0.0001$ and $\lambda = 0.1$ in GBM

| $k$ | Driver pathway | $W$ | $n_1$ | $n_2$ | $\lambda \cdot Z$ | $U$ | $P_W$ | $P_U$ |
|---|---|---|---|---|---|---|---|---|
| 2 | *TP53, CDKN2A*$^*_{mg}$ | 73 | 253 | 37 | 10.4111 | 83.4111 | 0 | 0 |
| 3 | *TP53, CDKN2A*$^*_{mg}$, *MDM2*$_{mg}$ | 79 | 250 | 47 | 10.8864 | 89.8864 | 0 | 0 |
| 4 | *CDK4*$_{mg}$, *CDKN2A*$^*_{mg}$, *RB1*$^{**}$, *OBSCN* | 83 | 181 | 35 | 8.8820 | 91.8820 | 0 | 0 |
| 5 | *CDK4*$_{mg}$, *CDKN2A*$^*_{mg}$, *RB1*$^{**}$, *OBSCN*, *MYCN*$_{mg}$ | 85 | 162 | 32 | 8.4232 | 93.4232 | 0 | 0 |
| 6 | *CDK4*$_{mg}$, *CDKN2A*$^*_{mg}$, *RB1*$^{**}$, *OBSCN*, *MYCN*$_{mg}$, *CDKN2A*$^{**}$ | 86 | 168 | 34 | 8.6385 | 94.6385 | 0 | 0 |
| 7 | *CDK4*$_{mg}$, *CDKN2A*$^*_{mg}$, *RB1*$^{**}$, *OBSCN*, *MYCN*$_{mg}$, *CDKN2A*$^{**}$, *RB1*$^*_{mg}$ | 86 | 190 | 39 | 9.2994 | 95.2994 | 0 | 0 |
| 8 | *CDK4*$_{mg}$, *CDKN2A*$^*_{mg}$, *RB1*$^{**}$, *OBSCN*, *MYCN*$_{mg}$, *CDKN2A*$^{**}$, *RB1*$^*_{mg}$, *B2M* | 87 | 169 | 36 | 8.8370 | 95.8370 | 0 | 0 |
| 9 | *CDK4*$_{mg}$, *CDKN2A*$^*_{mg}$, *RB1*$^{**}$, *OBSCN*, *MYCN*$_{mg}$, *CDKN2A*$^{**}$, *RB1*$^*_{mg}$, *B2M, PHLPP1* | 86 | 187 | 37 | 9.1973 | 95.1973 | 0 | 0 |
| 10 | *CDK4*$_{mg}$, *CDKN2A*$^*_{mg}$, *RB1*$^{**}$, *OBSCN*, *MYCN*$_{mg}$, *CDKN2A*$^{**}$, *RB1*$^*_{mg}$, *B2M, PHLPP1, GABRB2* | 85 | 193 | 41 | 9.5264 | 94.5264 | 0 | 0 |

$W$ represents the weight value defined by Eq. (**1**) for the identified driver pathway (gene set); $n_1$ and $n_2$ represent the numbers of impacted genes and TFs or miRNAs, respectively; $Z = Z_{S,E} + Z_{S,A}$ is a score reflecting the impacts of the gene set on the expression of downstream genes and the activities of TFs and miRNAs, where $Z_{S,E}$ and $Z_{S,A}$ are defined in Eqs. **4** and **5**, respectively; $U$ is the sum of $W$ and $\lambda \cdot Z$, denoting the new weight for driver pathway defined in Eq. **6**; $P_W$ and $P_U$ denote the $p$-values of the identified driver gene set for the weights $W$ and $U$, respectively.

- *CDKN2A*$^*_{mg}$ represents the deletion on chromosome region 9p21.3, including metagenes *CDKN2A, CDKN2B, CDKN2B-AS1, IFNA1, IFNA2, IFNA5, IFNA6, IFNA8, IFNA22P, IFNA13, MTAP, IFNE, DMRTA1, MIR31, KLHL9, C9orf53.*
- *CDKN2A*$^{**}$ denotes gene *CDKN2A* mutation.
- *RB1*$^*_{mg}$ represents the deletion on chromosome region 13q14.2, including metagenes *RB1, LPAR6.*
- *RB1*$^{**}$ denotes gene *RB1* mutation.
- *MDM2*$_{mg}$ denotes metagenes representing the amplification of *MDM2, LYZ, FRS2, CPM, YEATS4.*
- *CDK4*$_{mg}$ denotes metagenes representing the amplification of *CDK4, MARCH9, TSPAN31.*
- *MYCN*$_{mg}$ denotes metagenes representing the amplification of *MYCN, MYCNOS.*
- *SIRT3*$_{mg}$ denotes metagenes representing the deletion of *SIRT3, PKP3, SCGB1C1, ATHL1, IFITM2, ANO9, NLRP6, B4GALNT4, LOC100133161, SIGIRR, PSMD13, IFITM5, IFITM1, LOC653486, BET1L, RIC8A, ODF3, IFITM3, DEAF1, POLR2L, RNH1, LSP1, FAM99A, CTSD, MIR4298, TSPAN4, KRTAP5-6, SYT8, BRSK2, DRD4, OR51E1, DNHD1, TNNI2, TMEM80, FAM99B.*

### 2 The performance of the proposed method on randomly selected gene sets

To validate the effectiveness of gtePIDP, we investigate its performance on randomly selected gene sets in GBM. Without loss of generality, we take $k = 5$ as an example for the analysis, and the results of 10 runs are displayed in Table S13.

Note that for this study we simultaneously consider two kinds of significance, i.e., $P_W$-value and $P_U$-value. Now for all the 10 runs, $P_W$-value is larger than 0.05, so all the detected gene sets cannot be candidate driver gene sets although for the ninth run the $P_U$-value is less than 0.05. Here for the 10 runs the randomly selected gene sets have mutation frequency ranges of 0.01-0.34, 0.02-0.44, 0.01-0.14, 0.01-0.05, 0.01-0.05, 0.01-0.18, 0.01-0.05, 0.01-0.17, 0.01-0.61, and 0.01-0.16.

### 3 The regulatory module related to apoptosis in GBM

Other more complicated regulatory modules can also be identified and constructed by the proposed method (for example, the functional module related to apoptosis in GBM, Fig. S1), where candidate therapeutic axes also deserve to be explored and further studied.

**Table S2** Identified significant driver gene sets for $\beta = 0.001$ and $\lambda = 0.1$ in GBM

| $k$ | Driver pathway | $W$ | $n_1$ | $n_2$ | $\lambda \cdot Z$ | $U$ | $P_W$ | $P_U$ |
|---|---|---|---|---|---|---|---|---|
| 2 | $TP53$, $CDKN2A^*_{mg}$ | 73 | 348 | 67 | 11.5598 | 84.5598 | 0 | 0 |
| 3 | $TP53$, $CDKN2A^*_{mg}$, $MDM2_{mg}$ | 79 | 348 | 76 | 11.9336 | 90.9336 | 0 | 0 |
| 4 | $CDK4_{mg}$, $CDKN2A^*_{mg}$, $RB1^{**}$, $OBSCN$ | 83 | 280 | 62 | 10.1754 | 93.1754 | 0 | 0 |
| 5 | $CDK4_{mg}$, $CDKN2A^*_{mg}$, $RB1^{**}$, $OBSCN$, $MYCN_{mg}$ | 85 | 255 | 63 | 9.8095 | 94.8095 | 0 | 0 |
| 6 | $CDK4_{mg}$, $CDKN2A^*_{mg}$, $RB1^{**}$, $OBSCN$, $MYCN_{mg}$, $CDKN2A^{**}$ | 86 | 264 | 66 | 10.0080 | 96.0080 | 0 | 0 |
| 7 | $CDK4_{mg}$, $CDKN2A^*_{mg}$, $RB1^{**}$, $OBSCN$, $MYCN_{mg}$, $CDKN2A^{**}$, $RB1^*_{mg}$ | 86 | 288 | 77 | 10.6932 | 96.6932 | 0 | 0 |
| 8 | $CDK4_{mg}$, $CDKN2A^*_{mg}$, $RB1^{**}$, $OBSCN$, $MYCN_{mg}$, $CDKN2A^{**}$, $RB1^*_{mg}$, $B2M$ | 87 | 272 | 76 | 10.3647 | 97.3647 | 0 | 0 |
| 9 | $CDK4_{mg}$, $CDKN2A^*_{mg}$, $RB1^{**}$, $OBSCN$, $MYCN_{mg}$, $CDKN2A^{**}$, $RB1^*_{mg}$, $B2M$, $PHLPP1$ | 86 | 292 | 79 | 10.7230 | 96.7230 | 0 | 0 |
| 10 | $CDK4_{mg}$, $CDKN2A^*_{mg}$, $RB1^{**}$, $OBSCN$, $MYCN_{mg}$, $CDKN2A^{**}$, $RB1^*_{mg}$, $B2M$, $PHLPP1$, $GABRB2$ | 85 | 313 | 76 | 10.9823 | 95.9823 | 0 | 0 |

The symbols have same meanings as in Table S1.

**Table S3** Identified significant driver gene sets for $\beta = 0.005$ and $\lambda = 0.1$ in GBM

| $k$ | Driver pathway | $W$ | $n_1$ | $n_2$ | $\lambda \cdot Z$ | $U$ | $P_W$ | $P_U$ |
|---|---|---|---|---|---|---|---|---|
| 2 | $CDK4_{mg}$, $CDKN2A^*_{mg}$ | 75 | 354 | 83 | 10.3727 | 85.3727 | 0 | 0 |
| 3 | $CDK4_{mg}$, $CDKN2A^*_{mg}$, $RB1^{**}$ | 81 | 373 | 89 | 10.7354 | 91.7354 | 0 | 0 |
| 4 | $CDK4_{mg}$, $CDKN2A^*_{mg}$, $RB1^{**}$, $OBSCN$ | 83 | 374 | 95 | 10.9538 | 93.9538 | 0 | 0 |
| 5 | $CDK4_{mg}$, $CDKN2A^*_{mg}$, $RB1^{**}$, $OBSCN$, $MYCN_{mg}$ | 85 | 350 | 97 | 10.6595 | 95.6595 | 0 | 0 |
| 6 | $CDK4_{mg}$, $CDKN2A^*_{mg}$, $RB1^{**}$, $OBSCN$, $MYCN_{mg}$, $CDKN2A^{**}$ | 86 | 358 | 102 | 10.8233 | 96.8233 | 0 | 0 |
| 7 | $CDK4_{mg}$, $CDKN2A^*_{mg}$, $RB1^{**}$, $OBSCN$, $MYCN_{mg}$, $CDKN2A^{**}$, $B2M$ | 87 | 347 | 98 | 10.4952 | 97.4952 | 0 | 0 |
| 8 | $CDK4_{mg}$, $CDKN2A^*_{mg}$, $RB1^{**}$, $OBSCN$, $MYCN_{mg}$, $CDKN2A^{**}$, $B2M$, $RB1^*_{mg}$ | 87 | 383 | 102 | 11.1264 | 98.1264 | 0 | 0 |
| 9 | $CDK4_{mg}$, $CDKN2A^*_{mg}$, $RB1^{**}$, $OBSCN$, $MYCN_{mg}$, $CDKN2A^{**}$, $B2M$, $RB1^*_{mg}$, $PHLPP1$ | 86 | 399 | 106 | 11.4473 | 97.4473 | 0 | 0 |
| 10 | $CDK4_{mg}$, $CDKN2A^*_{mg}$, $ATM$, $OBSCN$, $CCND2_{mg}$, $CDKN2A^{**}$, $RB1^*_{mg}$, $B2M$, $PHLPP1$, $NOTCH1$ | 85 | 414 | 108 | 11.6826 | 96.6826 | 0 | 0 |

The symbols have same meanings as in Table S1. In addition, $CCND2_{mg}$ denotes metagenes representing the amplification of *CCND2, KDM5A, CCDC77, B4GALNT3, SLC6A13, IQSEC3, LOC574538, WNK1, SLC6A12, CCND2, LOC100288778, NINJ2, FAM138D, ADIPOR2.*

### 4 Application to other malignancies

#### 4.1 Application to BRCA and OV

For BRCA and OV, the identified driver gene sets are listed in Tables S14 and S15, respectively. As two typical gynecological tumors, previous studies manifested that there are some commonalities between them. Using the proposed method gtePIDP, we found that much more difference exists between these two cancers, especially exists for the impact of driver genes on TFs and miRNAs as well as their regulation on downstream gene expressions.

**Table S4** Identified significant driver gene sets for $\beta = 0.01$ and $\lambda = 0.1$ in GBM

| $k$ | Driver pathway | $W$ | $n_1$ | $n_2$ | $\lambda \cdot Z$ | $U$ | $P_W$ | $P_U$ |
|---|---|---|---|---|---|---|---|---|
| 2 | $CDK4_{mg}$, $CDKN2A^*_{mg}$ | 75 | 391 | 106 | 10.6705 | 85.6705 | 0 | 0 |
| 3 | $CDK4_{mg}$, $CDKN2A^*_{mg}$, $RB1^{**}$ | 81 | 421 | 108 | 11.0162 | 92.0162 | 0 | 0 |
| 4 | $CDK4_{mg}$, $CDKN2A^*_{mg}$, $RB1^{**}$, *OBSCN* | 83 | 430 | 110 | 11.2192 | 94.2192 | 0 | 0 |
| 5 | $CDK4_{mg}$, $CDKN2A^*_{mg}$, $RB1^{**}$, *OBSCN*, $MYCN_{mg}$ | 85 | 411 | 114 | 10.9702 | 95.9702 | 0 | 0 |
| 6 | $CDK4_{mg}$, $CDKN2A^*_{mg}$, $RB1^{**}$, *OBSCN*, $MYCN_{mg}$, $CDKN2A^{**}$ | 86 | 416 | 118 | 11.1058 | 97.1058 | 0 | 0 |
| 7 | $CDK4_{mg}$, $CDKN2A^*_{mg}$, $RB1^{**}$, *OBSCN*, $MYCN_{mg}$, $CDKN2A^{**}$, *B2M* | 87 | 404 | 111 | 10.7663 | 97.7663 | 0 | 0 |
| 8 | *TP53*, $CDKN2A^*_{mg}$, $MDM2_{mg}$, $CDKN2A^{**}$, *GABRA1*, *GABRB2*, $SIRT3_{mg}$, *PHLPP1* | 78 | 523 | 148 | 13.5609 | 91.5609 | 0 | 0 |
| 9 | $CDK4_{mg}$, $CDKN2A^*_{mg}$, $RB1^{**}$, *OBSCN*, $MYCN_{mg}$, $CDKN2A^{**}$, *B2M*, $RB1^*_{mg}$, *PHLPP1* | 86 | 451 | 123 | 11.6968 | 97.6968 | 0 | 0 |
| 10 | $CDK4_{mg}$, $CDKN2A^*_{mg}$, $RB1^{**}$, *OBSCN*, $MYCN_{mg}$, $CDKN2A^{**}$, *B2M*, $RB1^*_{mg}$, *PHLPP1*, *CACNA1A* | 85 | 460 | 129 | 11.9732 | 96.9732 | 0 | 0 |

The symbols have same meanings as in Table S1.

**Table S5** Identified significant driver gene sets for $\beta = 0.0001$ and $\lambda = 0.3$ in GBM

| $k$ | Driver pathway | $W$ | $n_1$ | $n_2$ | $\lambda \cdot Z$ | $U$ | $P_W$ | $P_U$ |
|---|---|---|---|---|---|---|---|---|
| 2 | *TP53*, $CDKN2A^*_{mg}$ | 73 | 253 | 37 | 31.2333 | 104.2333 | 0 | 0 |
| 3 | *TP53*, $CDKN2A^*_{mg}$, $MDM2_{mg}$ | 79 | 250 | 47 | 32.6592 | 111.6592 | 0 | 0 |
| 4 | *TP53*, $CDKN2A^*_{mg}$, $MDM2_{mg}$, $CDKN2A^{**}$ | 80 | 258 | 49 | 33.2897 | 113.2897 | 0 | 0 |
| 5 | *TP53*, $CDKN2A^*_{mg}$, $MDM2_{mg}$, $CDKN2A^{**}$, $RB1^*_{mg}$ | 80 | 266 | 51 | 34.0786 | 114.0786 | 0 | 0 |
| 6 | *TP53*, $CDKN2A^*_{mg}$, $MDM2_{mg}$, $CDKN2A^{**}$, $RB1^*_{mg}$, *GABRB2* | 79 | 281 | 55 | 35.3022 | 114.3022 | 0 | 0 |
| 7 | $CDK4_{mg}$, $CDKN2A^*_{mg}$, $RB1^{**}$, *OBSCN*, *CDKN2C*, $RB1^*_{mg}$, *RBP1* | 79 | 323 | 77 | 39.2499 | 118.2499 | 0 | 0 |
| 8 | $CDK4_{mg}$, $CDKN2A^*_{mg}$, $RB1^{**}$, *OBSCN*, *CDKN2C*, $RB1^*_{mg}$, *RBP1*, $CDKN2A^{**}$ | 80 | 334 | 75 | 39.6705 | 119.6705 | 0 | 0 |
| 9 | $CDK4_{mg}$, $CDKN2A^*_{mg}$, $RB1^{**}$, *OBSCN*, *CDKN2C*, $RB1^*_{mg}$, *RBP1*, $CDKN2A^{**}$, *PHLPP1* | 79 | 352 | 77 | 40.7938 | 119.7938 | 0 | 0 |
| 10 | *TP53*, $CDKN2A^*_{mg}$, $MDM2_{mg}$, $CDKN2A^{**}$, $RB1^*_{mg}$, *GABRB2*, *CACNA1A*, $SIRT3_{mg}$, *PHLPP1*, *PKD2* | 75 | 320 | 63 | 38.5037 | 113.5037 | 0 | 0 |

The symbols have same meanings as in Table S1.

#### 4.2 Application to LUAD

For LUAD, the identified driver gene sets are listed in Table S16. Take $k = 8$ as an example. We identified *TP53*, *STK11*, $KIF26B_{mg}$, $MDM2_{mg}$, *SETD2*, *MLL*, *CMAS*, $DLGAP1_{mg}$ as the driver gene set, which influenced activity changes of 142 TFs or miRNAs, and differential expressions of 389 downstream genes. Among these 389 genes there are 65 hallmark genes, six prognostic genes, and four cancer dependent genes.

We also found that there are a large proportion of downstream disturbed genes encoding glycoproteins or related to alternative splicing in LUAD (glycoproteins: 37.63%, alternative splicing: 43.04%). The disturbed biological processes or functions by the detected driver gene sets including cell migration, cell adhesion, inflammatory response, positive regulation of cell proliferation, negative regulation of cell differentiation, cell surface receptor linked signal transduction, enzyme linked receptor protein signaling pathway, etc.

**Table S6** Identified significant driver gene sets for $\beta = 0.001$ and $\lambda = 0.3$ in GBM

| $k$ | Driver pathway | $W$ | $n_1$ | $n_2$ | $\lambda \cdot Z$ | $U$ | $P_W$ | $P_U$ |
|---|---|---|---|---|---|---|---|---|
| 2 | *TP53*, $CDKN2A^*_{mg}$ | 73 | 348 | 67 | 34.6795 | 107.6795 | 0 | 0 |
| 3 | *TP53*, $CDKN2A^*_{mg}$, $MDM2_{mg}$ | 79 | 348 | 76 | 35.8008 | 114.8008 | 0 | 0 |
| 4 | *TP53*, $CDKN2A^*_{mg}$, $MDM2_{mg}$, $CDKN2A^{**}$ | 80 | 360 | 80 | 36.5140 | 116.5140 | 0 | 0 |
| 5 | *TP53*, $CDKN2A^*_{mg}$, $MDM2_{mg}$, $CDKN2A^{**}$, $RB1^*_{mg}$ | 80 | 366 | 84 | 37.2821 | 117.2821 | 0 | 0 |
| 6 | $CDK4_{mg}$, $CDKN2A^*_{mg}$, $RB1^{**}$, *CDKN2C*, *RBP1*, *OBSCN* | 79 | 420 | 104 | 40.6722 | 119.6722 | 0 | 0 |
| 7 | $CDK4_{mg}$, $CDKN2A^*_{mg}$, $RB1^{**}$, *CDKN2C*, *RBP1*, *OBSCN*, $RB1^*_{mg}$ | 79 | 439 | 107 | 42.1930 | 121.1930 | 0 | 0 |
| 8 | $CDK4_{mg}$, $CDKN2A^*_{mg}$, $RB1^{**}$, *CDKN2C*, *RBP1*, *OBSCN*, $RB1^*_{mg}$, $CDKN2A^{**}$ | 80 | 434 | 108 | 42.4467 | 122.4467 | 0 | 0 |
| 9 | $CDK4_{mg}$, $CDKN2A^*_{mg}$, $RB1^{**}$, *CDKN2C*, *RBP1*, *OBSCN*, $RB1^*_{mg}$, $CDKN2A^{**}$, *PHLPP1* | 79 | 454 | 113 | 43.5750 | 122.5750 | 0 | 0 |
| 10 | $CDK4_{mg}$, $CDKN2A^*_{mg}$, $RB1^{**}$, *CDKN2C*, *RBP1*, *OBSCN*, $RB1^*_{mg}$, $CDKN2A^{**}$, *PHLPP1*, *CACNA1A* | 78 | 472 | 121 | 44.6053 | 122.6053 | 0 | 0 |

The symbols have same meanings as in Table S1.

**Table S7** Identified significant driver gene sets for $\beta = 0.005$ and $\lambda = 0.3$ in GBM

| $k$ | Driver pathway | $W$ | $n_1$ | $n_2$ | $\lambda \cdot Z$ | $U$ | $P_W$ | $P_U$ |
|---|---|---|---|---|---|---|---|---|
| 2 | *TP53*, $CDKN2A^*_{mg}$ | 73 | 428 | 101 | 36.5532 | 109.5532 | 0 | 0 |
| 3 | *TP53*, $CDKN2A^*_{mg}$, $MDM2_{mg}$ | 79 | 445 | 103 | 37.5747 | 116.5747 | 0 | 0 |
| 4 | $CDK4_{mg}$, $CDKN2A^*_{mg}$, *RBP1*, $RB1^{**}$ | 78 | 480 | 119 | 39.0578 | 117.0578 | 0 | 0 |
| 5 | $CDK4_{mg}$, $CDKN2A^*_{mg}$, *RBP1*, $RB1^{**}$, *OBSCN* | 80 | 477 | 120 | 39.6336 | 119.6336 | 0 | 0 |
| 6 | $CDK4_{mg}$, $CDKN2A^*_{mg}$, *RBP1*, $RB1^{**}$, *OBSCN*, $RB1^*_{mg}$ | 80 | 508 | 131 | 41.3892 | 121.3892 | 0 | 0 |
| 7 | $CDK4_{mg}$, $CDKN2A^*_{mg}$, *RBP1*, $RB1^{**}$, *OBSCN*, $RB1^*_{mg}$, $CDKN2A^{**}$ | 81 | 510 | 133 | 41.7387 | 122.7387 | 0 | 0 |
| 8 | $CDK4_{mg}$, $CDKN2A^*_{mg}$, *RBP1*, $RB1^{**}$, *OBSCN*, $RB1^*_{mg}$, $CDKN2A^{**}$, *CDKN2C* | 80 | 534 | 147 | 44.0877 | 124.0877 | 0 | 0 |
| 9 | $CDK4_{mg}$, $CDKN2A^*_{mg}$, $RB1^*_{mg}$, *RBP1*, $RB1^{**}$, *OBSCN*, $CDKN2A^{**}$, *CDKN2C*, *CACNA1A* | 79 | 551 | 153 | 44.9955 | 123.9955 | 0 | 0 |
| 10 | $CDK4_{mg}$, $CDKN2A^*_{mg}$, $RB1^*_{mg}$, *RBP1*, $RB1^{**}$, *OBSCN*, $CDKN2A^{**}$, *CDKN2C*, *CACNA1A*, *PHLPP1* | 78 | 558 | 153 | 45.8737 | 123.8737 | 0 | 0 |

The symbols have same meanings as in Table S1.

**Table S8** Identified significant driver gene sets for $\beta = 0.01$ and $\lambda = 0.3$ in GBM

| $k$ | Driver pathway | $W$ | $n_1$ | $n_2$ | $\lambda \cdot Z$ | $U$ | $P_W$ | $P_U$ |
|---|---|---|---|---|---|---|---|---|
| 2 | $TP53,\ CDKN2A^*_{mg}$ | 73 | 481 | 119 | 37.2196 | 110.2196 | 0 | 0 |
| 3 | $TP53,\ CDKN2A^*_{mg},\ MDM2_{mg}$ | 79 | 495 | 122 | 38.2254 | 117.2254 | 0 | 0 |
| 4 | $TP53,\ CDKN2A^*_{mg},\ MDM2_{mg},\ CDKN2A^{**}$ | 80 | 498 | 125 | 38.7271 | 118.7271 | 0 | 0 |
| 5 | $TP53,\ CDKN2A^*_{mg},\ MDM2_{mg},\ CDKN2A^{**},\ RB1^*_{mg}$ | 80 | 503 | 126 | 39.3664 | 119.3664 | 0 | 0 |
| 6 | $CDK4_{mg},\ CDKN2A^*_{mg},\ OBSCN,\ RBP1,\ RB1^*_{mg},\ RB1^{**}$ | 80 | 554 | 151 | 41.9299 | 121.9299 | 0 | 0 |
| 7 | $CDK4_{mg},\ CDKN2A^*_{mg},\ CDKN2C,\ CDKN2A^{**},\ RB1^*_{mg},\ RB1^{**},\ NF1$ | 82 | 517 | 153 | 40.6385 | 122.6385 | 0 | 0 |
| 8 | $CDK4_{mg},\ CDKN2A^*_{mg},\ CDKN2C,\ CDKN2A^{**},\ RB1^*_{mg},\ RB1^{**},\ RBP1,\ OBSCN$ | 80 | 586 | 167 | 44.5641 | 124.5641 | 0 | 0 |
| 9 | $CDK4_{mg},\ CDKN2A^*_{mg},\ CDKN2C,\ CDKN2A^{**},\ RB1^*_{mg},\ RB1^{**},\ RBP1,\ OBSCN,\ PHLPP1$ | 79 | 596 | 169 | 45.4643 | 124.4643 | 0 | 0 |
| 10 | $CDK4_{mg},\ CDKN2A^*_{mg},\ CDKN2C,\ CDKN2A^{**},\ RB1^*_{mg},\ RB1^{**},\ RBP1,\ OBSCN,\ PHLPP1,\ CACNA1A$ | 78 | 598 | 173 | 46.2627 | 124.2627 | 0 | 0 |

The symbols have same meanings as in Table S1.

**Table S9** Identified significant driver gene sets for $\beta = 0.0001$ and $\lambda = 0.5$ in GBM

| $k$ | Driver pathway | $W$ | $n_1$ | $n_2$ | $\lambda \cdot Z$ | $U$ | $P_W$ | $P_U$ |
|---|---|---|---|---|---|---|---|---|
| 2 | $TP53,\ CDKN2A^*_{mg}$ | 73 | 253 | 37 | 52.0556 | 125.0556 | 0 | 0 |
| 3 | $TP53,\ CDKN2A^*_{mg},\ MDM2_{mg}$ | 79 | 250 | 47 | 54.4320 | 133.4320 | 0 | 0 |
| 4 | $TP53,\ CDKN2A^*_{mg},\ MDM2_{mg},\ CDKN2A^{**}$ | 80 | 258 | 49 | 55.4828 | 135.4828 | 0 | 0 |
| 5 | $TP53,\ CDKN2A^*_{mg},\ MDM2_{mg},\ CDKN2A^{**},\ CDKN2C$ | 77 | 295 | 56 | 59.8698 | 136.8698 | 0 | 0 |
| 6 | $CDK4_{mg},\ CDKN2A^*_{mg},\ CDKN2C,\ RB1^{**},\ RBP1,\ RB1^*_{mg}$ | 77 | 323 | 77 | 64.7302 | 141.7302 | 0 | 0 |
| 7 | $TP53,\ CDKN2A^*_{mg},\ MDM2_{mg},\ CDKN2A^{**},\ RBP1,\ RB1^*_{mg},\ GABRB2$ | 72 | 358 | 67 | 67.2318 | 139.2318 | 0 | 0 |
| 8 | $CDK4_{mg},\ CDKN2A^*_{mg},\ CDKN2C,\ RB1^{**},\ RBP1,\ RB1^*_{mg},\ CDKN2A^{**},\ OBSCN$ | 80 | 334 | 75 | 66.1175 | 146.1175 | 0 | 0 |
| 9 | $CDK4_{mg},\ CDKN2A^*_{mg},\ CDKN2C,\ RB1^{**},\ RBP1,\ RB1^*_{mg},\ CDKN2A^{**},\ OBSCN,\ PHLPP1$ | 79 | 352 | 77 | 67.9897 | 146.9897 | 0 | 0 |
| 10 | $CDK4_{mg},\ CDKN2A^*_{mg},\ CDKN2C,\ RB1^{**},\ RBP1,\ RB1^*_{mg},\ CDKN2A^{**},\ OBSCN,\ PHLPP1,\ GABRB2$ | 78 | 362 | 78 | 69.4378 | 147.4378 | 0 | 0 |

The symbols have same meanings as in Table S1.

**Table S10** Identified significant driver gene sets for $\beta = 0.001$ and $\lambda = 0.5$ in GBM

| $k$ | Driver pathway | $W$ | $n_1$ | $n_2$ | $\lambda \cdot Z$ | $U$ | $P_W$ | $P_U$ |
|---|---|---|---|---|---|---|---|---|
| 2 | $TP53,\ CDKN2A^*_{mg}$ | 73 | 348 | 67 | 57.7992 | 130.7992 | 0 | 0 |
| 3 | $TP53,\ CDKN2A^*_{mg},\ MDM2_{mg}$ | 79 | 348 | 76 | 59.6681 | 138.6681 | 0 | 0 |
| 4 | $TP53,\ CDKN2A^*_{mg},\ MDM2_{mg},\ CDKN2A^{**}$ | 80 | 360 | 80 | 60.8567 | 140.8567 | 0 | 0 |
| 5 | $CDK4_{mg},\ CDKN2A^*_{mg},\ RB1^{**},\ RBP1,\ CDKN2C$ | 77 | 401 | 100 | 66.4857 | 143.4857 | 0 | 0 |
| 6 | $CDK4_{mg},\ CDKN2A^*_{mg},\ RB1^{**},\ RBP1,\ CDKN2C,\ OBSCN$ | 79 | 420 | 104 | 67.7870 | 146.7870 | 0 | 0 |
| 7 | $TP53,\ CDKN2A^*_{mg},\ MDM2_{mg},\ CDKN2A^{**},\ RB1^*_{mg},\ RBP1,\ GABRB2$ | 72 | 469 | 109 | 72.5614 | 144.5614 | 0 | 0 |
| 8 | $CDK4_{mg},\ CDKN2A^*_{mg},\ RB1^*_{mg},\ CDKN2C,\ RBP1,\ RB1^{**},\ OBSCN,\ CDKN2A^{**}$ | 80 | 434 | 108 | 70.7445 | 150.7445 | 0 | 0 |
| 9 | $TP53,\ CDKN2A^*_{mg},\ MDM2_{mg},\ CDKN2A^{**},\ RB1^*_{mg},\ RBP1,\ GABRB2,\ PHLPP1,\ CACNA1A$ | 70 | 491 | 114 | 75.2381 | 145.2381 | 0 | 0 |
| 10 | $CDK4_{mg},\ CDKN2A^*_{mg},\ RB1^*_{mg},\ CDKN2C,\ RBP1,\ RB1^{**},\ OBSCN,\ CDKN2A^{**},\ PHLPP1,\ CACNA1A$ | 78 | 472 | 121 | 74.3422 | 152.3422 | 0 | 0 |

The symbols have same meanings as in Table S1.

**Table S11** Identified significant driver gene sets for $\beta = 0.005$ and $\lambda = 0.5$ in GBM

| $k$ | Driver pathway | $W$ | $n_1$ | $n_2$ | $\lambda \cdot Z$ | $U$ | $P_W$ | $P_U$ |
|---|---|---|---|---|---|---|---|---|
| 2 | $TP53,\ CDKN2A^*_{mg}$ | 73 | 428 | 101 | 60.9219 | 133.9219 | 0 | 0 |
| 3 | $TP53,\ CDKN2A^*_{mg},\ MDM2_{mg}$ | 79 | 445 | 103 | 62.6245 | 141.6245 | 0 | 0 |
| 4 | $TP53,\ CDKN2A^*_{mg},\ MDM2_{mg},\ CDKN2A^{**}$ | 80 | 448 | 106 | 63.5351 | 143.5351 | 0 | 0 |
| 5 | $CDK4_{mg},\ CDKN2A^*_{mg},\ RBP1,\ RB1^{**},\ CDKN2C$ | 77 | 502 | 133 | 69.2362 | 146.2362 | 0 | 0 |
| 6 | $CDK4_{mg},\ CDKN2A^*_{mg},\ RBP1,\ RB1^{**},\ CDKN2C,\ OBSCN$ | 79 | 511 | 138 | 70.3210 | 149.3210 | 0 | 0 |
| 7 | $TP53,\ CDKN2A^*_{mg},\ MDM2_{mg},\ CDKN2A^{**},\ CDKN2C,\ RB1^*_{mg},\ GABRB2$ | 76 | 513 | 130 | 70.8535 | 146.8535 | 0 | 0 |
| 8 | $CDK4_{mg},\ CDKN2A^*_{mg},\ RBP1,\ RB1^{**},\ CDKN2C,\ OBSCN,\ CDKN2A^{**},\ RB1^*_{mg}$ | 80 | 534 | 147 | 73.4794 | 153.4794 | 0 | 0 |
| 9 | $TP53,\ CDKN2A^*_{mg},\ MDM2_{mg},\ CDKN2A^{**},\ CDKN2C,\ RB1^*_{mg},\ GABRB2,\ SIRT3_{mg},\ MLLT4_{mg}$ | 72 | 558 | 154 | 75.5473 | 147.5473 | 0 | 0 |
| 10 | $CDK4_{mg},\ CDKN2A^*_{mg},\ RBP1,\ RB1^{**},\ CDKN2C,\ OBSCN,\ CDKN2A^{**},\ RB1^*_{mg},\ PHLPP1,\ CACNA1A$ | 78 | 558 | 153 | 76.4562 | 154.4562 | 0 | 0 |

The symbols have same meanings as in Table S1. In addition, $MLLT4_{mg}$ denotes metagenes representing the deletion of *MLLT4, QKI, UNC93A, TCP10, GPR31, TCP10L2, PARK2, TTLL2, C6orf123.*

**Table S12** Identified significant driver gene sets for $\beta = 0.01$ and $\lambda = 0.5$ in GBM

| $k$ | Driver pathway | $W$ | $n_1$ | $n_2$ | $\lambda \cdot Z$ | $U$ | $P_W$ | $P_U$ |
|---|---|---|---|---|---|---|---|---|
| 2 | $TP53,\ CDKN2A^*_{mg}$ | 73 | 481 | 119 | 62.0327 | 135.0327 | 0 | 0 |
| 3 | $TP53,\ CDKN2A^*_{mg},\ MDM2_{mg}$ | 79 | 495 | 122 | 63.7090 | 142.7090 | 0 | 0 |
| 4 | $TP53,\ CDKN2A^*_{mg},\ MDM2_{mg},\ CDKN2A^{**}$ | 80 | 498 | 125 | 64.5452 | 144.5452 | 0 | 0 |
| 5 | $CDK4_{mg},\ CDKN2A^*_{mg},\ RBP1,\ RB1^{**},\ CDKN2C$ | 77 | 547 | 156 | 70.1881 | 147.1881 | 0 | 0 |
| 6 | $CDK4_{mg},\ CDKN2A^*_{mg},\ RBP1,\ RB1^{**},\ CDKN2C,\ OBSCN$ | 79 | 555 | 157 | 71.1270 | 150.1270 | 0 | 0 |
| 7 | $CDK4_{mg},\ CDKN2A^*_{mg},\ RBP1,\ RB1^{**},\ CDKN2C,\ OBSCN,\ RB1^*_{mg}$ | 79 | 577 | 168 | 73.7545 | 152.7545 | 0 | 0 |
| 8 | $TP53,\ CDKN2A^*_{mg},\ MDM2_{mg},\ CDKN2A^{**},\ CDKN2C,\ RB1^*_{mg},\ GABRB2,\ SIRT3_{mg}$ | 75 | 575 | 158 | 73.1375 | 148.1375 | 0 | 0 |
| 9 | $CDK4_{mg},\ CDKN2A^*_{mg},\ RBP1,\ RB1^{**},\ CDKN2C,\ OBSCN,\ RB1^*_{mg},\ CDKN2A^{**},\ PHLPP1$ | 79 | 596 | 169 | 75.7739 | 154.7739 | 0 | 0 |
| 10 | $CDK4_{mg},\ CDKN2A^*_{mg},\ RBP1,\ RB1^{**},\ CDKN2C,\ OBSCN,\ RB1^*_{mg},\ CDKN2A^{**},\ PHLPP1,\ CACNA1A$ | 78 | 598 | 173 | 77.1045 | 155.1045 | 0 | 0 |

The symbols have same meanings as in Table S1.

**Table S13** Randomly selected gene sets for $k = 5$ in GBM

| Driver pathway | $W$ | $n_1$ | $n_2$ | $\lambda \cdot Z$ | $U$ | $P_W$ | $P_U$ |
|---|---|---|---|---|---|---|---|
| $BRCA2_{mg},\ TP53,\ PRKG1_{mg},\ GABRB2,\ KRAS$ | 36 | 3 | 0 | 2.0740 | 38.0740 | 0.6900 | 0.6900 |
| $LPA,\ PTEN^{**},\ CCND2_{mg},\ EGFR^*_{mg},\ PIK3R1$ | 51 | 16 | 3 | 7 | 58 | 0.1400 | 0.0500 |
| $PTEN^*_{mg},\ OBSCN,\ GABRA1,\ PDGFRA_{mg},\ PIK3CA^{**}$ | 26 | 3 | 0 | 2.0698 | 28.0698 | 0.3500 | 0.3900 |
| $PIK3CA^{**},\ ATM,\ PIK3CA^*_{mg},\ PTPRD_{mg},\ CCDC144A$ | 10 | 0 | 0 | 0 | 10 | 0.9000 | 0.9300 |
| $PTPRD_{mg},\ FBXW7_{mg},\ CDKN2C,\ NEFH,\ SETD2$ | 10 | 0 | 0 | 0 | 10 | 0.9800 | 0.9900 |
| $GABRA1,\ GTPBP2_{mg},\ EGFR^{**},\ MYC_{mg},\ MLLT4_{mg}$ | 20 | 0 | 0 | 0 | 20 | 0.9600 | 0.9000 |
| $KIF13B,\ FBXW7_{mg},\ GSTP1,\ LPA,\ NOTCH1$ | 11 | 0 | 0 | 0 | 11 | 0.8700 | 0.9000 |
| $NOTCH1,\ PTEN^{**},\ CDK4_{mg},\ PHLPP1,\ GSTP1$ | 30 | 2 | 0 | 1.7421 | 31.7421 | 0.7000 | 0.6200 |
| $CDKN2A^*_{mg},\ PHLPP1,\ NF1,\ B2M,\ GABRB2$ | 63 | 95 | 9 | 17 | 80 | 0.1500 | 0.0200 |
| $MET,\ MYCN_{mg},\ CDK4_{mg},\ KRAS,\ CDK6$ | 23 | 0 | 0 | 0 | 23 | 0.2700 | 0.2300 |

The symbols have same meanings as in Table S1. In addition, $BRCA2_{mg}$ denotes the metagenes including *BRCA2, ITGAL, POLE, ZEB2, PAN3, KIAA0368, GATA3, PHF6, TNXB, CABLES1, CDKN1B, MAPKBP1, TAF4, MYO1B, EPHA2, CUL1, FAM5C, TRIM7, NRG1, NOTCH2, PDE8B*; $PRKG1_{mg}$ denotes the metagenes including *PRKG1, MIR605, CSTF2T*; $PTEN^*_{mg}$ represents the deletion on chromosome region 10q23.31, including metagenes *PTEN, KLLN*; $PTEN^{**}$ denotes gene *PTEN* mutation; $CCND2_{mg}$ denotes the metagenes including *CCND2, C12orf5, KDM5A, CCDC77, B4GALNT3, SLC6A13, IQSEC3, LOC574538, WNK1, SLC6A12, LOC100288778, NINJ2, FAM138D, ADIPOR2*; $EGFR^*_{mg}$ represents the amplification on chromosome region 7p11.2, including metagenes *EGFR, VOPP1*; $EGFR^{**}$ denotes gene *EGFR* mutation; $PDGFRA_{mg}$ denotes the metagenes including *PDGFRA, PDCL2, KIT, NMU*; $PIK3CA^*_{mg}$ represents the amplification on chromosome region 3q26, including metagenes *PIK3CA, SOX2, MFN1, TERC, GNB4, MECOM, ZMAT3, KCNMB3, ZNF639, KCNMB2*; $PIK3CA^{**}$ denotes gene *PIK3CA* mutation; $PTPRD_{mg}$ denotes the metagenes including *PTPRD, C9orf66, DOCK8, FOXD4, RFX3, FAM138C, WASH1, CBWD1, KANK1*; $FBXW7_{mg}$ denotes the metagenes including *FBXW7, PLCL1*; $\mathrm{GTPBP2}_{mg}$ denotes the metagenes including *GTPBP2, MRPS18A, TJAP1, CCND3, TBCC, PPP2R5D, MAD2L1BP, PEX6, GNMT, PTCRA, RPL7L1, YIPF3, PRPH2, POLR1C, POLH, RSPH9, MEA1, C6orf226, KIAA0240, CNPY3, LOC100132354, KLHDC3, CUL7, UBR2, XPO5, C6orf223, VEGFA*; $MYC_{mg}$ denotes the metagenes including *MYC, POU5F1B, LOC727677, MIR1204, MIR1208, MIR1205, PVT1*; $MLLT4_{mg}$ denotes the metagenes including *MLLT4, QKI, UNC93A, TCP10, GPR31, TCP10L2, PARK2, TTLL2, C6orf123.*

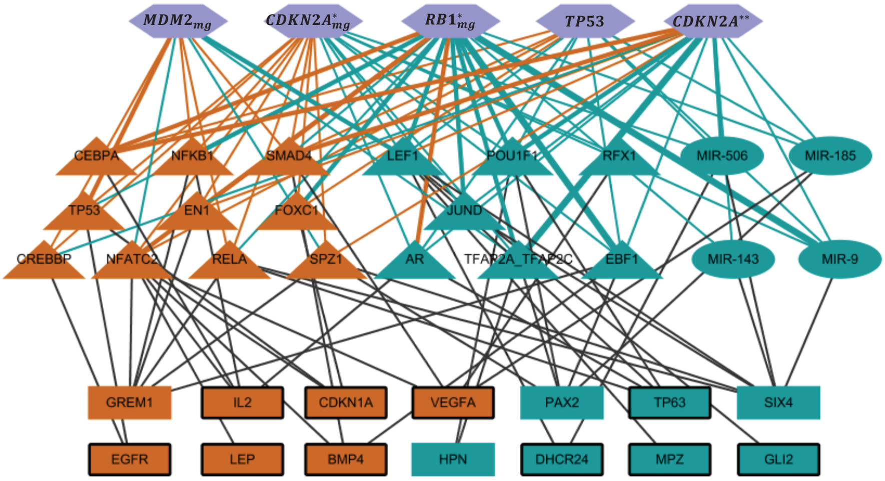


**Fig. S1** The regulatory module related to apoptosis in GBM.

**Table S14** Identified significant driver gene sets for $\beta = 0.0001$ and $\lambda = 0.3$ in BRCA

| $k$ | Driver pathway | $W$ | $n_1$ | $n_2$ | $\lambda \cdot Z$ | $U$ | $P_W$ | $P_U$ |
|---|---|---|---|---|---|---|---|---|
| 2 | *TP53, PIK3CA* | 236 | 68 | 4 | 13.1484 | 249.1484 | 0 | 0 |
| 3 | *TP53, PIK3CA, GATA3* | 260 | 69 | 6 | 14.1343 | 274.1343 | 0 | 0 |
| 4 | *TP53, PIK3CA, GATA3, CDH1* | 272 | 117 | 11 | 19.2906 | 291.2906 | 0 | 0 |
| 5 | *TP53, PIK3CA, GATA3, CDH1, CTCF* | 279 | 121 | 10 | 19.1548 | 298.1548 | 0 | 0 |
| 6 | *TP53, PIK3CA, GATA3, CDH1, CTCF,* $FLRT3_{mg}$ | 283 | 128 | 14 | 20.6584 | 303.6584 | 0 | 0 |
| 7 | *TP53, PIK3CA, GATA3, CDH1, CTCF,* $FLRT3_{mg}$, $MYCN_{mg}$ | 281 | 167 | 32 | 25.6908 | 306.6908 | 0 | 0 |
| 8 | *TP53, PIK3CA, GATA3, CDH1, CTCF,* $FLRT3_{mg}$, $MYCN_{mg}$, *MAPK1* | 283 | 176 | 32 | 26.3281 | 309.3281 | 0 | 0 |
| 9 | *TP53, PIK3CA, GATA3, CDH1, CTCF,* $FLRT3_{mg}$, $MYCN_{mg}$, *MAPK1, PIK3R1* | 285 | 182 | 33 | 26.7007 | 311.7007 | 0 | 0 |
| 10 | *TP53, PIK3CA, GATA3, CDH1, NF1,* $FLRT3_{mg}$, *LPA, AKT1, PIK3R1, ZZEF1* | 288 | 169 | 31 | 25.7821 | 313.7821 | 0 | 0 |

The symbols have same meanings as in Table S1. In addition, $FLRT3_{mg}$ denotes metagenes representing the deletion of *FLRT3, MACROD2*.

**Table S15** Identified significant driver gene sets for $\beta = 0.0001$ and $\lambda = 0.3$ in OV

| $k$ | Driver pathway | $W$ | $n_1$ | $n_2$ | $\lambda \cdot Z$ | $U$ | $P_W$ | $P_U$ |
|---|---|---|---|---|---|---|---|---|
| 2 | *TP53, $WNT2B_{mg}$* | 230 | 543 | 101 | 51.6623 | 281.6623 | 0.01 | 0 |
| 3 | *TP53, $WNT2B_{mg}$, KRAS* | 231 | 541 | 103 | 51.8009 | 282.8009 | 0 | 0 |
| 4 | *TP53, $WNT2B_{mg}$, KRAS, NRAS* | 231 | 548 | 99 | 51.8982 | 282.8982 | 0 | 0 |
| 5 | *TP53, $WNT2B_{mg}$, KRAS, NRAS, FAM124B* | 230 | 555 | 102 | 52.3578 | 282.3578 | 0 | 0 |
| 6 | *TP53, $WNT2B_{mg}$, KRAS, NRAS, FAM124B, CDK17* | 229 | 556 | 104 | 52.5522 | 281.5522 | 0 | 0 |
| 7 | *TP53, $WNT2B_{mg}$, KRAS, NRAS, FAM124B, CDK17, FLT3* | 228 | 557 | 105 | 52.6333 | 280.6333 | 0 | 0 |
| 8 | *TP53, $WNT2B_{mg}$, KRAS, NRAS, FAM124B, CDK17, FLT3, MAP3K1* | 227 | 556 | 105 | 52.6729 | 279.6729 | 0 | 0 |
| 9 | *TP53, $WNT2B_{mg}$, KRAS, NRAS, FAM124B, CDK17, FBXW7, MAP3K1, MAP2K4* | 226 | 556 | 105 | 52.6838 | 278.6838 | 0 | 0 |
| 10 | *TP53, $WNT2B_{mg}$, KRAS, NRAS, FAM124B, CDK17, FBXW7, MAP3K1, MAP2K4, KCTD1* | 225 | 559 | 106 | 52.7189 | 277.7189 | 0 | 0 |

The symbols have same meanings as in Table S1. In addition, $WNT2B_{mg}$ denotes metagenes representing the deletion of *WNT2B, TRIM33, SYT6, MAB21L3, ATP1A1OS*.

**Table S16** Identified significant driver gene sets for $\beta = 0.0001$ and $\lambda = 0.3$ in LUAD

| $k$ | Driver pathway | $W$ | $n_1$ | $n_2$ | $\lambda \cdot Z$ | $U$ | $P_W$ | $P_U$ |
|---|---|---|---|---|---|---|---|---|
| 2 | *TP53, KRAS* | 130 | 300 | 70 | 38.6244 | 168.6244 | 0 | 0 |
| 3 | *TP53, KRAS, $KIF26B_{mg}$* | 126 | 388 | 128 | 48.4823 | 174.4823 | 0.0100 | 0 |
| 4 | *TP53, STK11, $KIF26B_{mg}$, $MDM2_{mg}$* | 131 | 378 | 129 | 48.5867 | 179.5867 | 0 | 0 |
| 5 | *TP53, STK11, $KIF26B_{mg}$, $MDM2_{mg}$, SETD2* | 135 | 381 | 133 | 48.8225 | 183.8225 | 0 | 0 |
| 6 | *TP53, STK11, $KIF26B_{mg}$, $MDM2_{mg}$, SETD2, MLL* | 136 | 397 | 148 | 50.5006 | 186.5006 | 0 | 0 |
| 7 | *TP53, STK11, $KIF26B_{mg}$, $MDM2_{mg}$, SETD2, MLL, CMAS* | 137 | 401 | 150 | 50.9939 | 187.9939 | 0 | 0 |
| 8 | *TP53, STK11, $KIF26B_{mg}$, $MDM2_{mg}$, SETD2, MLL, CMAS, $DLGAP1_{mg}$* | 139 | 389 | 142 | 50.2309 | 189.2309 | 0 | 0 |
| 9 | *TP53, $CDK4_{mg}$, $MDM4_{mg}$, ATM, SETD2, PIK3R1, RBM10, PFN2, $RPL5_{mg}$* | 144 | 407 | 132 | 49.2580 | 193.2580 | 0 | 0 |
| 10 | *TP53, STK11, $KIF26B_{mg}$, $MDM2_{mg}$, SETD2, MLL, CMAS, $DLGAP1_{mg}$, MAGEL2, PFN2* | 141 | 396 | 149 | 51.0021 | 192.0021 | 0 | 0 |

The symbols have same meanings as in Table S1. In addition, $KIF26B_{mg}$ denotes metagenes representing the amplification of *KIF26B, C1orf100, C1orf31, C1orf150, ACTN2*; $DLGAP1_{mg}$ denotes metagenes representing the deletion of *DLGAP1, FLJ35776, LOC284215, LOC201477*; $MDM4_{mg}$ denotes metagenes representing the amplification of *MDM4, PIK3C2B, NFASC, LOC127841, REN, GOLT1A, ETNK2, PPP1R15B, KISS1, PLEKHA6, SOX13, CNTN2, LRRN2, C1orf157*; $RPL5_{mg}$ denotes metagenes representing the deletion of *RPL5, SNORA66, SNORD21*.
